# *Physcomitrium patens* SMXL homologs are PpMAX2-dependent negative regulators of growth

**DOI:** 10.1101/2023.04.11.536386

**Authors:** Ambre Guillory, Mauricio Lopez-Obando, Khalissa Bouchenine, Louis Lambret, Philippe Le Bris, Alain Lécureuil, Jean-Paul Pillot, Vincent Steinmetz, François-Didier Boyer, Catherine Rameau, Alexandre de Saint Germain, Sandrine Bonhomme

## Abstract

SMXL proteins are a plant-specific clade of type I HSP100/Clp-ATPases. *SMXL* genes are found in virtually all land plant genomes. However, they have mainly been studied in angiosperms. In *Arabidopsis thaliana*, three SMXL functional subclades have been identified: SMAX1/SMXL2, SMXL345 and SMXL678. Out of these, two subclades ensure transduction of endogenous hormone signals: SMAX1/SMXL2 are involved in KAI2-ligand (KL) signaling, while SMXL678 are involved in strigolactone (SL) signaling. Many questions remain regarding the mode of action of these proteins, as well as their ancestral role. We addressed these questions by investigating the function of the four *SMXL* genes of the moss *Physcomitrium patens.* We demonstrate that PpSMXL proteins are involved in the conserved ancestral MAX2-dependent KL signaling pathway and act as negative regulators of growth. However, PpSMXL proteins expressed in *A. thaliana* unexpectedly cannot replace SMAX1/SMXL2 function in KL signaling, whereas they can functionally replace SMXL4/5 and restore root growth. Therefore, the molecular function of SMXL could be conserved, but not their interaction network. Moreover, one PpSMXL clade positively regulates transduction of the SL signal in *P. patens.* So far, this function has only been reported herein in moss, where it represents a novel crosstalk between SL and KL signaling pathways.

## Introduction

Strigolactones (SLs) are butenolide compounds with an early origin in land plant evolution (Delaux et al. 2012; Kyozuka et al. 2022; Bonhomme and Guillory 2022). In angiosperms, these molecules have first been identified as rhizospheric signals with both negative and positive outcomes for the producing plant: SLs stimulate seed germination of root parasitic plants (Cook et al. 1966) but also promote Arbuscular Mycorrhizal (AM) symbiosis (Akiyama et al. 2005) by boosting AM fungi mitochondrial metabolism and thus hyphae growth (Besserer et al. 2006; Besserer et al. 2008). SLs are also employed as phytohormones in angiosperms, where they play diverse roles, in particular regulating plant architecture (for review see (Machin et al. 2020)). Notably, SLs have a largely documented ability to repress axillary branching, by inhibiting axillary bud activity (Gomez-Roldan et al. 2008; Umehara et al. 2008; Jiang et al. 2013; Kerr et al. 2021).

Phylogenetic studies suggest that the SL biosynthesis pathway is ancient, with genes encoding most SL biosynthesis hormones being found both in land plants and some algae (Delaux et al. 2012; Walker et al. 2019). The CAROTENOID CLEAVAGE DIOXYGENASE 8 (CCD8) enzyme is essential for SL biosynthesis since its product, carlactone (CL), is considered a key precursor for SLs (Alder et al. 2012). Genetic studies have demonstrated that CCD8 function, first established in angiosperm models, is probably conserved in bryophytes, as this function was demonstrated at least in the moss *Physcomitrium patens* (*P. patens*) (Proust et al. 2011; Decker et al. 2017) and in the liverwort *Marchantia paleacea* (Radhakrishnan et al. 2020; Kodama et al. 2022). However, this conservation has not been proven yet for algal CCD8 homologs (Walker et al. 2019) and so far, SL synthesis has only been described in land plants. In *P. patens*, SLs are used as hormonal signals: Indeed, PpCCD8-derived compounds repress filament branching and elongation (Proust et al. 2011; Hoffmann et al. 2014), repress gametophore (leafy shoot) basal branching (Coudert et al. 2015), and enhance resistance to phytopathogenic fungi (Decker et al. 2017). Still, the exact molecule(s) with a hormonal role has(have) not been identified yet (Lopez-Obando et al. 2021). In contrast, in *M. paleacea*, the recently characterized SL Bryosymbiol (BSB) has no apparent hormonal function but is an essential rhizospheric signal needed for AM symbiosis establishment (Kodama et al. 2022).

In angiosperms, SL perception is achieved by the receptor DWARF14 (D14). SL perception triggers D14’s binding to the F-box protein MORE AXILLARY GROWTH 2 (MAX2), and subsequent targeting of repressor proteins for degradation by the 26S proteasome (Bennett et al. 2016). These repressors are encoded by a small gene family of type I HSP100/Clp-ATPases and are called SUPPRESSOR OF MAX2 (SMAX)1-LIKE (SMXL) in Arabidopsis (Stanga et al. 2013; Soundappan et al. 2015; Wang et al. 2015), and DWARF53 (D53) in rice (Jiang et al. 2013). SMXL6, 7, and 8 are key repressors of the SL pathway for branching control (Soundappan et al. 2015; Wang et al. 2015). SMAX1 and SMXL2 are repressors in a parallel signaling pathway controlling seed germination and seedling development in Arabidopsis (Stanga et al. 2016; Villaecija-Aguilar et al. 2019; Park et al. 2022), and mesocotyl elongation and AM symbiosis in rice (Zheng et al. 2020; Choi et al. 2020). This parallel pathway also involves the MAX2 F-box protein, and the KARRIKIN INSENSITIVE 2 (KAI2) receptor (Conn and Nelson 2016), a paralog of D14. While the existence of an endogenous hormonal KAI2-ligand (KL) is expected (Waters et al. 2015; Conn and Nelson 2016; Sun et al. 2016), the identity of molecule(s) activating the so-called KL pathway is still unknown.

Thanks to reverse genetics, SL and KL signaling pathways have been partially described in bryophyte models (Proust et al. 2011; Decker et al. 2017; Mizuno et al. 2021; Kodama et al. 2022; Kyozuka et al. 2022; Bonhomme and Guillory 2022). In *Marchantia polymorpha*, a KAI2 and MAX2-dependent pathway regulates thallus growth using a sole MpSMXL repressor (Mizuno et al. 2021). In the moss *P. patens*, we have previously shown that the homolog of MAX2 is not necessary for response to SLs but is involved in the response to red light and possibly in KL signaling (Lopez-Obando et al. 2018; Lopez-Obando et al. 2021). The *KAI2* gene family is extended in *P. patens*, comprising candidate receptors for SLs (GJM clade), and candidate receptors for KL (euKAI2 clade). Thus, both SL (MAX2-independent) and KL (MAX2-dependent) pathways are present in *P. patens*, in contrast to *M. polymorpha* and *M. paleacea* (Lopez-Obando et al. 2021, Kodama et al. 2022). Also contrasting with the single SMXL copy of *M. polymorpha,* there are four *PpSMXL* genes. Hence, we have sought to determine possible roles played by *P. patens* SMXL proteins in the ancestral MAX2-dependent KL signaling pathway and in the MAX2-independent, moss-specific SL signaling pathway.

Here, we show that the four PpSMXL proteins all display canonical SMXL domains and motifs (Walker et al. 2019) and collectively play a negative role downstream of PpMAX2 in the putative KL signaling pathway of *P. patens*. Introduction of moss *PpSMXL* genes in Arabidopsis failed to induce functional complementation of the *smax1* mutant, suggesting PpSMXL proteins are not able to form a functional signaling complex with AtKAI2 and AtMAX2. Our results nonetheless support the hypothesis of the ancestral role of SMXL proteins being in KL response. Moreover, two PpSMXL proteins could act as positive actors of SL signaling and thus constitute a level of crosstalk between the SL and the PpMAX2-dependent KL pathway. The finding that none of these SMXL homologs are repressors in the SL signaling pathway supports the hypothesis that the mechanism triggering SL signal transduction has an independent origin in *P. patens* and in seed plants, even though the same family of receptors was recruited in both lineages (Bythell-Douglas et al. 2017; Lopez-Obando et al. 2021). Furthermore, our work supports the hypothesis that the use of SLs as hormones is not an ancestral land plant trait (Kodama et al. 2022) and might not be shared by seedless plants outside of bryopsid mosses.

## Results

### Phylogeny and expression analysis reveal two clades of *SMXL* genes in *P. patens*

SMXL are large proteins with several conserved structural domains: a double Clp-N domain in the N terminus and two ATPase domains (D1 and D2), separated by a long middle region (M). The D2 domain is split into D2a and D2b subdomains by an Ethylene-responsive-element-binding factor Amphiphilic Repression (EAR) motif (for review see (Temmerman et al. 2022)). SMXL activity partly relies on the EAR motif mediating transcriptional repression (Liang et al. 2016; Ma et al. 2017). In the D2a subdomain lies a specific degron motif (RGKT or Walker-A P loop), necessary for KL/SL-triggered SMXL degradation (Zhou et al. 2013; Khosla et al. 2020).

The four PpSMXL proteins harbor this canonical SMXL domain organization (Supplemental Figure S1). As per a previous extensive analysis of SMXL proteins phylogeny (Walker et al. 2019), the PpSMXLA and PpSMXLB proteins belong to a SMXL clade that is specific to bryopsid mosses, while PpSMXLC and PpSMXLD correspond to a SMXL clade that is common to all mosses. At the amino acid level, PpSMXLC and PpSMXLD show 72% identity and PpSMXLA and PpSMXLB 61%, whereas comparisons between proteins of the two different clades give 27-29% identity.

We investigated *PpSMXL* genes’ expression along vegetative development by RT-qPCR and observed that transcript levels are generally lower in the *PpSMXLA/B* clade than in the *PpSMXLC/D* clade (Supplemental Figure S2)*. PpSMXL* transcript levels were mostly stable across protonema development, and tended to be higher in mature gametophores and/or rhizoids at later developmental stages (Supplemental Figure S2).

### *Ppsmxl* loss-of-function mutants do not display a constitutive SL response phenotype

To investigate the function of the four *PpSMXL* genes in *P. patens* and their potential involvement in the SL and KL pathways, we employed CRISPR-Cas9-mediated mutagenesis to yield either mutants with frameshift or small insertions/deletions (*Ppsmxl*) (Lopez-Obando et al. 2016b), or mutants with complete deletion of the CDS (*Ppsmxl*Δ) (Supplemental Figure S3 and S4). Reasoning that the four PpSMXL proteins grouping into two clades could reflect different functions (Supplemental Figure S2 and (Walker et al. 2019)), we primarily characterized double mutants for each clade, e.g. *Ppsmxl(*Δ*)ab* and *Ppsmxl(*Δ*)cd* mutants.

Single and double mutants in the *PpSMXLA/B* clade were similar to the WT when grown under white light (Figure 1, A-D). On the other hand, both *Ppsmxlcd* and *Ppsmxl*Δ*cd* double mutants showed markedly enhanced protonema extension, resulting in very large plants (Figure 1, A-D), even surpassing the *Ppccd8* mutant in some experiments (Figure 1D). Single *Ppsmxld* mutant also tended to be larger than WT, although not to the same level as double mutants (Figure 1, A-B). Thus, it appears *PpSMXLC/D* loss of function results in a protonema extension phenotype similar to *Ppccd8* and opposite to the *Ppmax2-1* mutant. The similarity between *Ppccd8* and *Ppsmxlcd* mutants was also clear when looked more closely at their protonema, both mutants displaying significantly increased protonema branching compared to WT (Supplemental Figure S5 A-C). A triple mutant (*Ppsmxla1b1d1*) and a quadruple mutant (*Ppsmxla2b2c1d2*) were also obtained. It should be noted that the *Ppsmxlc1* mutation in the quadruple mutant did not induce a frameshift but removed four residues in the D1 domain of the PpSMXLC protein (Supplemental Figure S4). When grown under white light, both multiple mutants displayed highly restrained growth and browning of the protonema, and only rare, stunted gametophores were produced (Figure 1 C-D). The dramatic phenotype of the quadruple mutant suggests that *PpSMXLA/B* and *PpSMXLC/D* clades may have redundant and essential functions for both protonema and gametophore development. Not exclusively, it may also indicate some kind of genetic interaction between both clades.

**Figure 1.**
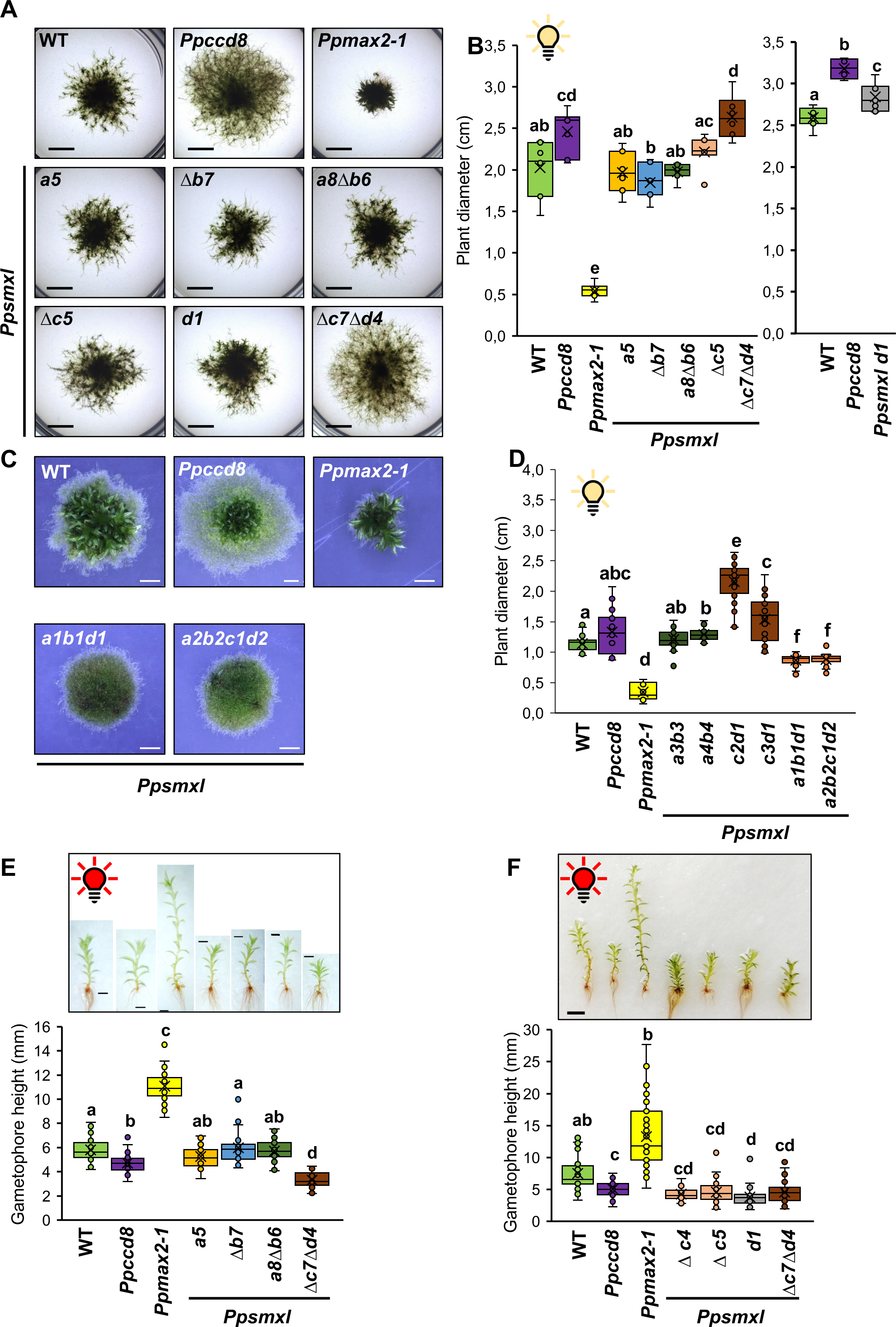
Phenotypes of the *Ppsmxl* mutants. *Ppsmxl* mutants were grown in white light long day conditions (A-D), or under continuous red light (E-F). A, Three-week-old plants grown in the light on low nitrogen content medium, without underlying cellophane. Bar = 5 mm. B, Diameters of six-week-old plants; Left panel: n=42-49 plants of each genotype grown on 6-7 individual plates. Statistical significance of comparisons between all genotypes are indicated by bold letters (Kruskal Wallis test (p<0.0001) followed by a Dunn *posthoc* test for pairwise comparisons); Right panel: n=20-35 plants of each genotype grown on at least 4 individual plates. Statistical significance of comparisons between all genotypes are indicated by bold symbols; right panel: standard ANOVA (p<0.0001) followed by a Tukey *post-hoc* test). C, Phenotype of two-week-old plants grown on low nitrogen content medium (with underlying cellophane). Scale bar = 2 mm. D, Diameters of 5-week-old plants; n=9-36 plants of each genotype, grown on 4 individual plates. Statistical significance of pairwise comparisons is indicated with letters (Welch’s ANOVA followed by a Dunnett’s T3 post-hoc test, p<0.05). E and F, Gametophores from two-month-old plants grown in red light (upper panels). Bar = 3 mm. Gametophore height was measured for n⩾30 two-month-old gametophores of each genotype (bottom panels). Statistical significance of comparisons between all genotypes are indicated by bold letters (Kruskal Wallis followed by a Dunn *post-hoc* test, p<5%).

We then assayed the phenotypes of the *Ppsmxl* mutants in conditions that previously allowed us to clearly distinguish mutants affected in the SL pathway from those affected in the PpMAX2-dependent KL pathway (Lopez-Obando et al. 2018; Lopez-Obando et al. 2021). Compared to WT, SL deficient (*Ppccd8*) and SL insensitive (*Ppkai2Lgjm*) mutants elongate more caulonema filaments in the dark, whereas KL insensitive mutants (*Ppmax2* and *eu-KAI2* clade) elongate fewer filaments in the dark and display a defective photomorphogenesis response to red light (Lopez-Obando et al. 2021). In the dark, *PpSMXLA/B* clade mutants were again alike WT, while *PpSMXLC/D* clade mutants developed more and longer filaments than WT, similarly to *Ppccd8* (Supplemental Figures S5 D-E). Puzzlingly, higher order *Ppsmxl* mutants grew few agravitropic curled caulonema filaments in the dark (Supplemental Figure S5 D-E). Under continuous red light, *PpSMXLA/B* clade mutants showed a similar phenotype to WT, while *PpSMXLC/D* clade mutants had smaller and more stunted gametophores, opposite to the *Ppmax2-1* mutant (Figure 1, E-F). Indeed, phytomer length was very short in *PpSMXLC/D* clade mutants (Supplemental Figure S6), suggesting constitutive photomorphogenesis.

Overall, *Ppsmxl* mutant phenotypic characterization suggests that (1) *PpSMXLC* and *PpSMXLD* act redundantly to limit protonema growth, at least partly by limiting branching of caulonema filaments, thereby regulating filament number and length; (2) *PpSMXLC* and *PpSMXLD* repress gametophore photomorphogenesis under red light; (3) *PpSMXLA* and *PpSMXLB* play a very modest role (if any) in regulation of protonema and gametophore growth in various light conditions.

### *PpSMXL* overexpressing lines display phenotypes alike *Ppmax2-1*

To further explore the putative role of PpSMXL as repressors of the PpMAX2-dependent pathway, we examined the phenotype of stable *P. patens* transgenic lines overexpressing *PpSMXLA* and *PpSMXLC* with a N-terminal GFP tag under the maize ubiquitin promoter (*proZmUbi:GFP-PpSMXL* lines, noted as OE-SMXLA and OE-SMXLC, Figure 2). Under standard growth conditions, both OE-SMXL lines were significantly less radially extended than the WT and developed fewer (but bigger) gametophores than WT (Figure 2, A-C). These lines also grew very elongated gametophores under continuous red-light (Figure 2, D-E), again behaving like *Ppmax2-1* and opposite to mutants of the *PpSMXLC/D* clade (comparing Figure 1, E-F to Figure 2, D-E). These features point to *PpSMXLC* but also *PpSMXLA* acting downstream of *PpMAX2* and playing an opposite role to *PpMAX2* in development.

**Figure 2.**
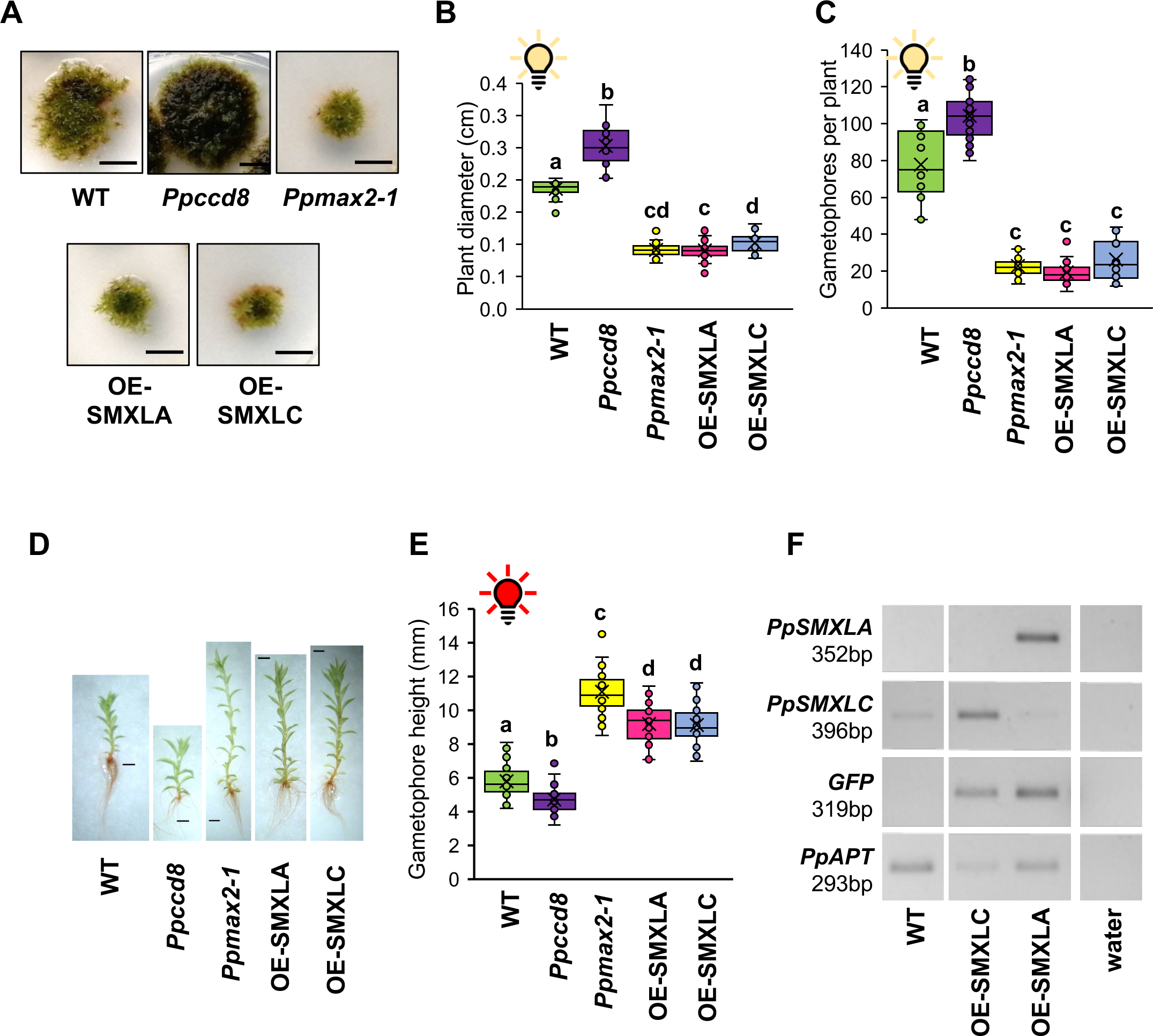
Growth of *proZmUbi:GFP-PpSMXL* lines. (A-B): Plant extension in control conditions. A, Representative individuals after 5 weeks of growth. Scale bar = 1 cm. B, Diameters were measured from 6 week-old plants, grown on at least 3 different plates. Statistical significance of comparisons between all genotypes are indicated as bold letters (Welch ANOVA (p<0.0001) followed by a Dunnett post-hoc test, n=21-35). C, Gametophore number was assessed on six-week-old plants, grown on at least 3 different plates. Statistical significance of of the comparison between all genotypes is indicated by bold letters (Welch’s ANOVA (p<0.0001) followed by a Dunnett’s T3 *post-hoc* test, n=14-21). (D-E) Gametophore height in red light. D, Representative individuals after two months culture under continuous red light. Scale bar = 1 mm. E, Gametophore height was measured on two-month-old gametophores of each genotype. Statistical significance of comparisons between all genotypes are indicated by bold letters (standard ANOVA (p<0.0001) followed by a Tukey *post-hoc* test, n=28-30). F, Expression of *GFP-PpSMXL* fusion transcripts in transgenic *P. patens* lines, as assessed by RT-PCR.

### *PpSMXL* loss-of-function does not restore WT growth in the *Ppccd8* background

To test for genetic links between *PpSMXL* genes and the SL pathway, we generated and characterized *Ppccd8 Ppsmxlab*, *Ppccd8 Ppsmxlcd*, and *Ppccd8 Ppsmxlabcd* mutants. We noticed that mutation of neither *PpSMXL* clade restored protonema extension to WT levels (Supplemental Figure S7). Strikingly, the mutation of all four *PpSMXL* genes, which had a dramatic negative effect on growth in the WT background (Figure 1, C-D and Supplemental Figure S5), was completely circumvented by the *Ppccd8* mutation, as the *Ppccd8 Ppsmxlabcd* quintuple mutant showed a phenotype similar to *Ppccd8*. (Supplemental Figure S7). It is however worthy of note that the *Ppsmxlc9* mutation in the *Ppccd8 Ppsmxlabcd* mutant does not lead to a frameshift, similar to the *Ppsmxlc1* mutation (Supplemental Figure S4).

These observations confirm that PpSMXL proteins are not acting as repressors in the SL pathway. More unexpectedly, the lack of functional PpSMXLA, B and D proteins appears to be detrimental only when endogenous SLs are present.

### *PpSMXL* mutations partially restore *Ppmax2* mutant phenotypes

To confirm the postulated genetic relationship between *PpSMXL* and *PpMAX2*, we tested for complementation of the *Ppmax2* mutant phenotype by *Ppsmxl* loss of function. For this, we mutated *PpSMXLA* and *PpSMXLB* in the *Ppmax2-1* mutant background, and mutated *PpMAX2* in one of the *Ppsmxl*Δ*cd* mutants (Supplemental Figure S8). We found that *Ppsmxlab* mutations did not improve *Ppmax2* protonema extension, while *Ppsmxlcd* mutations partially suppressed this mutant phenotype under standard growth conditions (Figure 3, A-B). However, both *Ppsmxlab* and *Ppsmxlcd* mutations could partially restore the excessive elongation of *Ppmax2* mutant gametophores under red light (Figure 3, C-D). Thus, all PpSMXL proteins could act as repressors of the PpMAX2-dependent KL pathway. Genes from *PpSMXLC/D* clade have a predominant role under standard growth conditions, while both *PpSMXL* clades seem to act redundantly in the regulation of gametophore elongation in red light.

**Figure 3.**
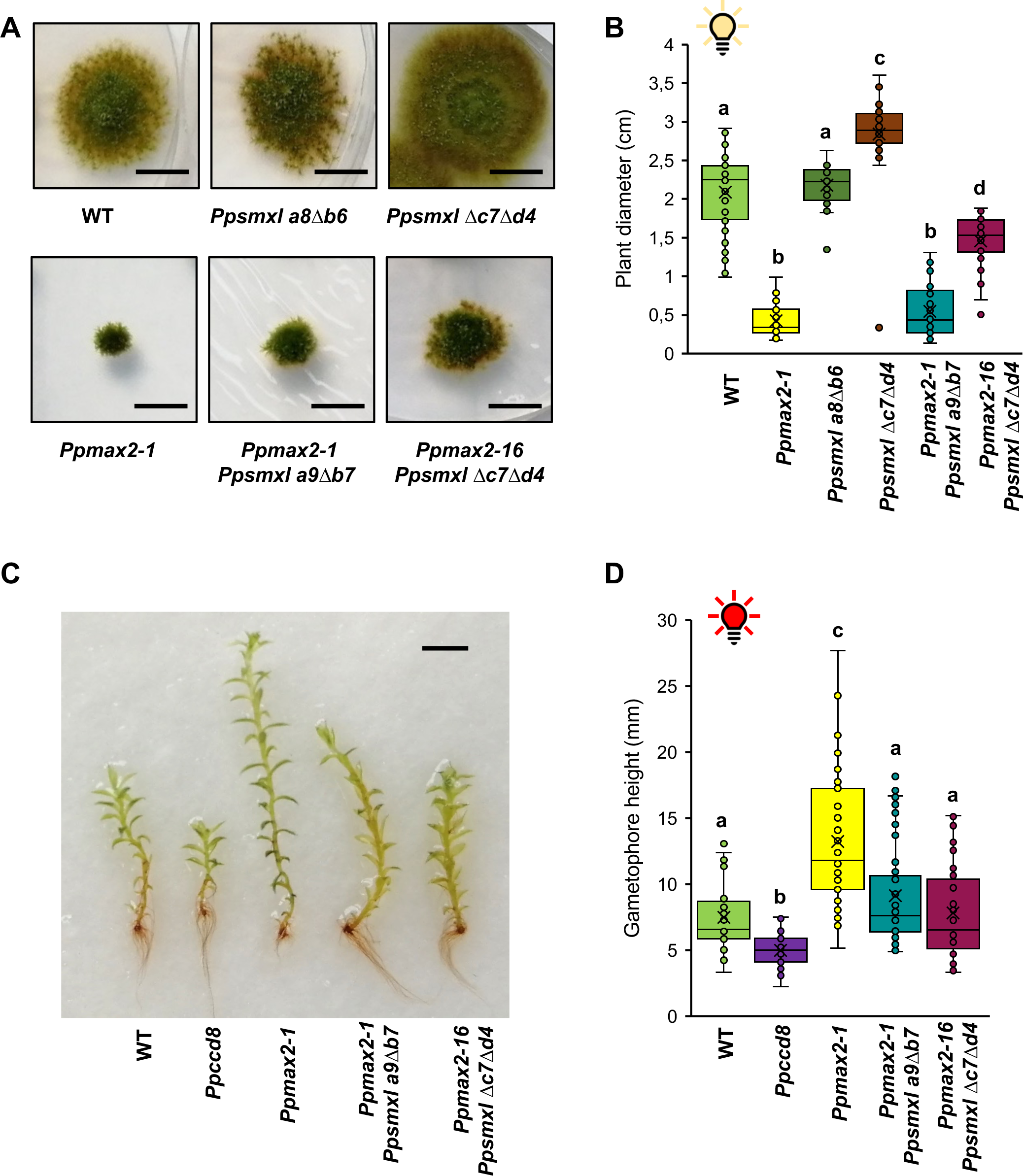
Genetic analysis of *PpSMXL* relationship with *PpMAX2*. (A-B): Plant extension in control conditions. A, Representative individuals after 5 weeks of growth. Bar = 1 cm. B, Plant diameters of the mutants grown on low nitrogen content medium with underlying cellophane for 5 weeks, with 7-8 independent plates per genotype. (C-D) Gametophore height in red light. C, Representative gametophores after two months culture under constant red light. Scale bar = 3 mm. D, Gametophore height was measured for 30 two-month-old gametophores of each genotype, with 3 independent Magenta boxes per genotype. (B,D): Statistical significance of comparisons between all genotypes are indicated by bold letters (Kruskal Wallis test (p<0.0001) followed by a Dunn *post-hoc* test, B: n=35-40; D: n=43-45).

### Loss of PpSMXL C/D function dampens the response to SL mimic but not to KL mimic

We then explored whether loss of *PpSMXL* function could impair the response to exogenously supplied SL and KL mimics. To this end, we assessed filament growth in the dark after application of GR24 enantiomers (Hoffmann et al. 2014; Lopez-Obando et al. 2021).

As the use of (−)-GR24 or KAR_2_ treatment to trigger the KL pathway previously gave non-reproducible results (Lopez-Obando et al. 2021), we instead chose to evaluate the effect of (−)-desmethyl-GR24 ((−)-dGR24), described as a better mimic for KL in both Arabidopsis and *Marchantia* (Yao et al. 2021). Increasing doses of (−)-dGR24 (0.01 to 10 µM) significantly increased the number of filaments in WT and *Ppccd8*, but had no effect on *Ppmax2-1*, as expected (Figure 4, A). On the other hand, the response to (−)-dGR24 was decreased but not abolished in the eu-KAI2 clade mutant, affected in *PpKAI2L-A* to *E* genes encoding putative KL receptors (Figure 4, A). Testing (−)-dGR24 on *Ppsmxl* double mutants, we observed that all of them were able to respond to this compound (here tested at 1 and 10 µM, Figure 4B). Considering four independent assays, the increase of filament number in response to 1 µM (−)-dGR24 was similar to WT for *Ppsmxlab* double mutant (Supplemental Figure S9A), whereas response was approximately two-fold lower for the *Ppsmxlcd* double mutant compared to WT (14.2 % versus 29.5 %, Supplemental Figure S9A). This observation leads to the hypothesis that *PpSMXLC/D* play a predominant role in response to (−)-desmethyl-GR24 in the dark.

**Figure 4.**
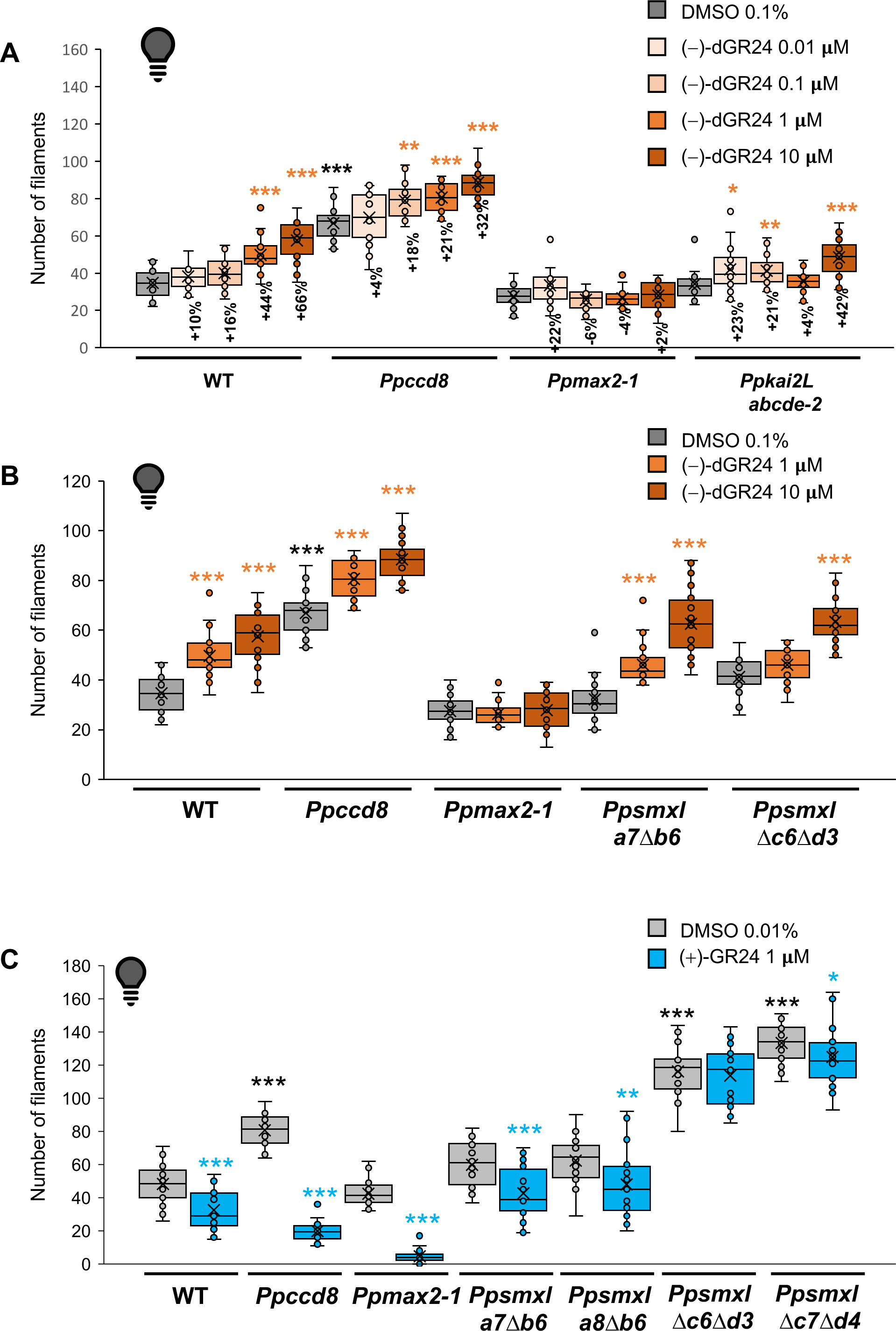
*Ppsmxl* mutant response to (+)-GR24 and (−)-desmethyl-GR24. Treatments were applied on two-week-old plants which were incubated vertically in the dark for ten days. Negatively gravitropic caulonema filaments were enumerated for each plant. A, Dose-response assays. plants were mock treated with 0,1% DMSO (grey) or increasing concentrations of (−)-desmethyl-GR24 ((−)-dGR24, light orange to brown : A, 0.01, 0.1, 1 and 10 µM). The gain or loss of filaments response has been calculated for each (−)-dGR24-treated group as the following ratio: (number_treated_ - number_mock_) / number_mock_. n=24 except for WT-DMSO and WT-0.01 µM where n=16. B, Response of *Ppsmxl* double mutants to (−)-dGR24: n=24 except for WT-DMSO; plants were mock treated with 0,1% DMSO (grey), 1 µM of (−)-dGR24 (orange) or 10 µM of (−)-dGR24 (brown). A and B, Statistical significance of comparisons between control and treated for each genotype is shown as bold orange symbols (Kruskal Wallis followed by a Dunn *post-hoc* test). Statistical significance of comparisons of control groups relative to WT is shown as bold black symbols (Kruskal Wallis test (p<0.0001) followed by a Dunn *post-hoc* test, n=16-24). C, Response to (+)-GR24: n = 24 two-week-old plants of each genotype were mock treated with 0,01% DMSO (grey) or with 1 µM of (+)-GR24 (blue). Statistical significance of comparisons between control and treated for each genotype is shown as blue symbols (Mann-Whitney tests). Statistical significance of comparisons of control groups relative to WT is shown as black symbols (Kruskal Wallis followed by a Dunn *post-hoc* test). For all statistical analyses, p-values are reported as * 0.01≤p<0.05, ** 0.001≤p<0.01, *** p<0.001.

When treated with 1 µM (+)-GR24 (SL mimic), WT and *Ppccd8* plants developed significantly less caulonema filaments, displaying a typical phenotypic response to SL treatment ((Lopez-Obando et al. 2021) and Figure 4, C). *Ppsmxl* single mutants and *Ppsmxlab* double mutants could respond to the SL mimic, apparently as much as the WT (Figure 4, C; Supplemental Figure S9, B-C). On the other hand, the response of *Ppsmxlcd* double mutants to (+)-GR24 was generally decreased compared to WT, considering five independent assays (Figure 4, C; Supplemental Figure S9, D). *PpSMXLC* and *PpSMXLD* play redundant roles in responding to (+)-GR24, as the response was not disturbed in single *Ppsmxlc* and *Ppsmxld* mutants (Supplemental Figure S9, C). This suggests that PpSMXLC/D could mediate at least part of the SL response, in addition to their role in the KL signaling.

### *PpSMXL* transcript levels are downregulated by light and modulated in response to both the SL and the KL signaling pathways

We wondered if a differential transcriptional regulation of *PpSMXL* genes could explain the different responses of moss mutants to SL and KL mimics. We therefore compared the transcript levels of *PpSMXL* genes in WT, *Ppccd8* and *Ppmax2-1* backgrounds. As the transcriptional response of SL-related genes is strongly affected by light and enhanced in the dark (Lopez-Obando et al. 2016a; Lopez-Obando et al. 2018), we assessed *PpSMXL* expression in both dark and light conditions.

We observed that all *PpSMXL* genes consistently had much higher transcript levels in the dark than in the light (Supplemental Figure S10, comparing the y-axis in left *versus* middle panels). In light conditions, *PpSMXLC* and *D* transcript levels were higher in *Ppccd8* compared to WT, indicating that endogenous SLs repress the expression of these two genes. Accordingly, (+)-GR24 application decreased *PpSMXLC* and *D* transcript levels, in the *Ppccd8* mutant (Supplemental Figure S10, C-D, right panels). Down-regulation of *PpSMXLC* and *PpSMXLD* transcripts by SLs further points to a possible crosstalk between the KL and SL pathways.

In the *Ppmax2-1* mutant, where PpSMXL proteins are likely over-accumulated, all *PpSMXL* genes were downregulated compared to WT in the dark (Supplemental Figure S10, A-D, left panels). This decrease was observed in the light as well, for *PpSMXLC* and *PpSMXLD* (Supplemental Figure S10, C-D, middle panels). This suggests that activation of the KL pathway upregulates the expression of *PpSMXL* genes.

Considered together, these results show that *PpSMXL* genes’ transcript levels are mainly down regulated by light and that expression of *PpSMXLC/D* genes responds to SL and KL signaling in opposite ways.

### PpSMXL proteins are localized mainly in the nucleus, and likely sensitive to a KL mimic but not to SL application

A sequence enriched in basic amino acids, corresponding to a functional N-terminal nuclear localization signal (NLS) reported in other SMXL proteins (Liang et al. 2016; Khosla et al. 2020; Choi et al. 2020), was found in the four PpSMXL proteins, suggesting a nuclear localization (Supplemental Figure S11). By transiently overexpressing RFP-PpSMXL fusion proteins in *Nicotiana benthamiana,* we could observe a clear nuclear signal for each PpSMXL protein, as evidenced by colocalization with the H2b-CFP histone marker (Supplemental Figure S12, A, C, E and G). Some cytosolic signal was also observed for PpSMXLA, PpSMXLB and PpSMXLC fusion proteins.

In angiosperms, the RGKT degron motif is responsible for SMXL proteasomal degradation, which is stimulated by SL or KL signaling (Zhou et al. 2013; Khosla et al. 2020). This motif is present in PpSMXLC and PpSMXLD proteins, and slightly modified in PpSMXLA and PpSMXLB (RGRT, Supplemental Figure S1; Supplemental Figure S12, I). In *N. benthamiana,* we found similarly high levels of nuclear RFP-PpSMXL fusion proteins whether the RGKT/RGRT motif was present or deleted (Supplemental Figure S12 B, D, F and H). Nonetheless, the additional cytosolic RFP signal seemed increased in leaves expressing ΔRGKT/RGRT variants, which could indicate an increase in RFP-PpSMXL stability (Supplemental Figure S12, D and H). Therefore, as for other SMXL proteins, PpSMXL turnover could depend on this degron motif.

Using our stable OE-PpSMXL *P. patens* lines, we could confirm the nuclear localization of PpSMXLA, PpSMXLC and PpSMXLD, even though the GFP signal was not always restricted to this compartment (Figure 5, A-B, and Supplemental Figure S13, A). The GFP-PpSMXLB fusion led to a faint and more diffuse cytoplasmic signal (Supplemental Figure S13, A). GFP-PpSMXLA fluorescent nuclei were easier to spot in the *Ppmax2-1* mutant background, suggesting PpMAX2 somehow affects the nuclear localization of the PpSMXLA protein (Figure 5, C). To further test the hypothesis of PpSMXL degradation through the KL pathway, we tested the effect of the KL mimic (−)-dGR24 on both OE-SMXLA and OE-SMXLC lines: a 20-minute-long treatment with 10 µM (−)-dGR24 decreased the nuclear signal, and the observed fluorescence was more diffuse and cytoplasmic, for both lines (Figure 5, A-B). The treatment with the proteasome inhibitor MG132 had no striking effect, even when applied concomitantly to (−)-dGR24. These observations suggest that PpSMXLA and PpSMXLC proteins are sensitive to a KL mimic.

**Figure 5.**
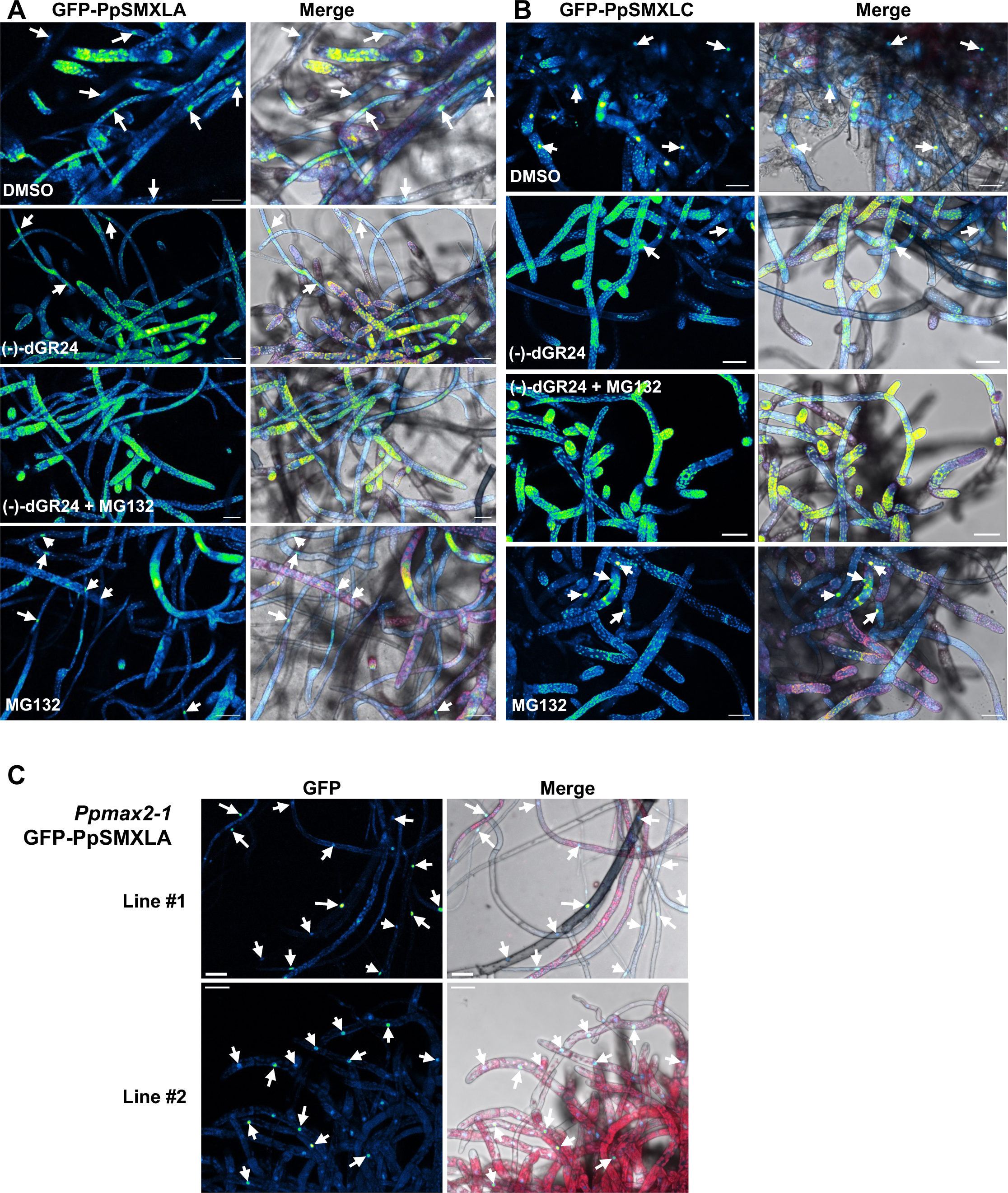
Subcellular localization of GFP-PpSMXLA and GFP-PpSMXLC fusion proteins in transgenic *P. patens* line and effect of (−)-dGR24 addition on stability. A, Localization of the GFP-PpSMXLA fusion protein in the WT background (OE-SMXLA line). B, Localization of the GFP-PpSMXLC fusion protein in the WT background (OE-SMXLC line). C, Localization of the GFP-PpSMXLA fusion protein in the *Ppmax2-1* background. Filaments were incubated in a solution of DMSO 0.01% (control) or in (-)-dGR24 at 10 µM (diluted in 0.01% DMSO) for 20 minutes, and/or treated with 100µM MG132 for the same duration. Merge are overlays of GFP (Green Fire Blue LUT), autofluorescence (Red) and bright field images. Scale bar = 50 µm. Instances of nuclei displaying GFP fluorescence are pointed at by white arrows.

We also tested a 20-minute-long treatment with 10 µM (+)-GR24 on the OE-SMXLA line, and observed no effect on the GFP-PpSMXLA signal, indicating that at least PpSMXLA is not rapidly degraded in response to SL (Supplemental Figure S13, B). Finally, fluorescent nuclei were also observed in *N. benthamiana* overexpressing the RFP-PpSMXL fusion proteins following a 5 µM (+)-GR24 treatment (Supplemental Figure S14). These results support the hypothesis that both PpSMXLA and PpSMXLC are sensitive to PpMAX2-dependent destabilization and are not degraded in response to SL.

### PpSMXL proteins can interact with players of the PpMAX2-dependent pathway

Since we hypothesized the PpSMXL proteins to act in the PpMAX2-dependent KL pathway, we sought to explore PpMAX2/PpSMXL, PpSMXL/PpKAI2L and PpKAI2L/PpMAX2 interactions. Using AlphaFold2-derived and ColabFold modeling (Jumper et al. 2021; Mirdita et al. 2021; Pettersen et al. 2021), we generated unbiased structure model of PpSMXLs - PpMAX2, PpSMXLs – PpKAI2L-C and PpMAX2 – PpKAI2L-C complexes. The models show large regions with predicted local Distance Difference Test (lDDT) scores <50, which are indicative of disordered regions in the PpSMXL proteins (Supplemental Figure S15, A) but PpMAX2, PpKAI2L-C, and D1, D2 and N domains of PpSMXLs were modeled with very high confidence. AlphaFold2 predicted that PpSMXLA and PpSMXLB could interact with PpMAX2 via their D2 domains with low predicted aligned error (PAE) values. In addition, we observed a high interaction probability between all PpSMXL D1 domains and the α-helices of PpKAI2L-C lid (Figure 6, A and Supplemental Figure S15, B), and between PpMAX2 LRR18 and 19 and the α-helices of PpKAI2L-C lid (Figure 6, A). To test these predictions *in planta*, we performed Bimolecular fluorescence complementation (BiFC) assays in *N. benthamiana.* BiFC suggested that each PpSMXL protein interacts with PpMAX2 in the nucleus (Figure 6, B, Supplemental Figures S15 and S16). Regarding PpSMXL/PpKAI2L interactions, we tested at least one PpKAI2L from each clade in the BiFC assays, including putative SL and KL receptors (Lopez-Obando et al. 2021). Only the putative KL receptor PpKAI2L-C from the eu-KAI2 clade could interact with the four PpSMXL proteins, also in the nucleus (Figure 6, B). To conclude, BiFC assays and *in silico* AlphaFold2 predictions both revealed interactions involving PpSMXL, PpMAX2 and a putative KL receptor, which would occur specifically in the nucleus.

**Figure 6.**
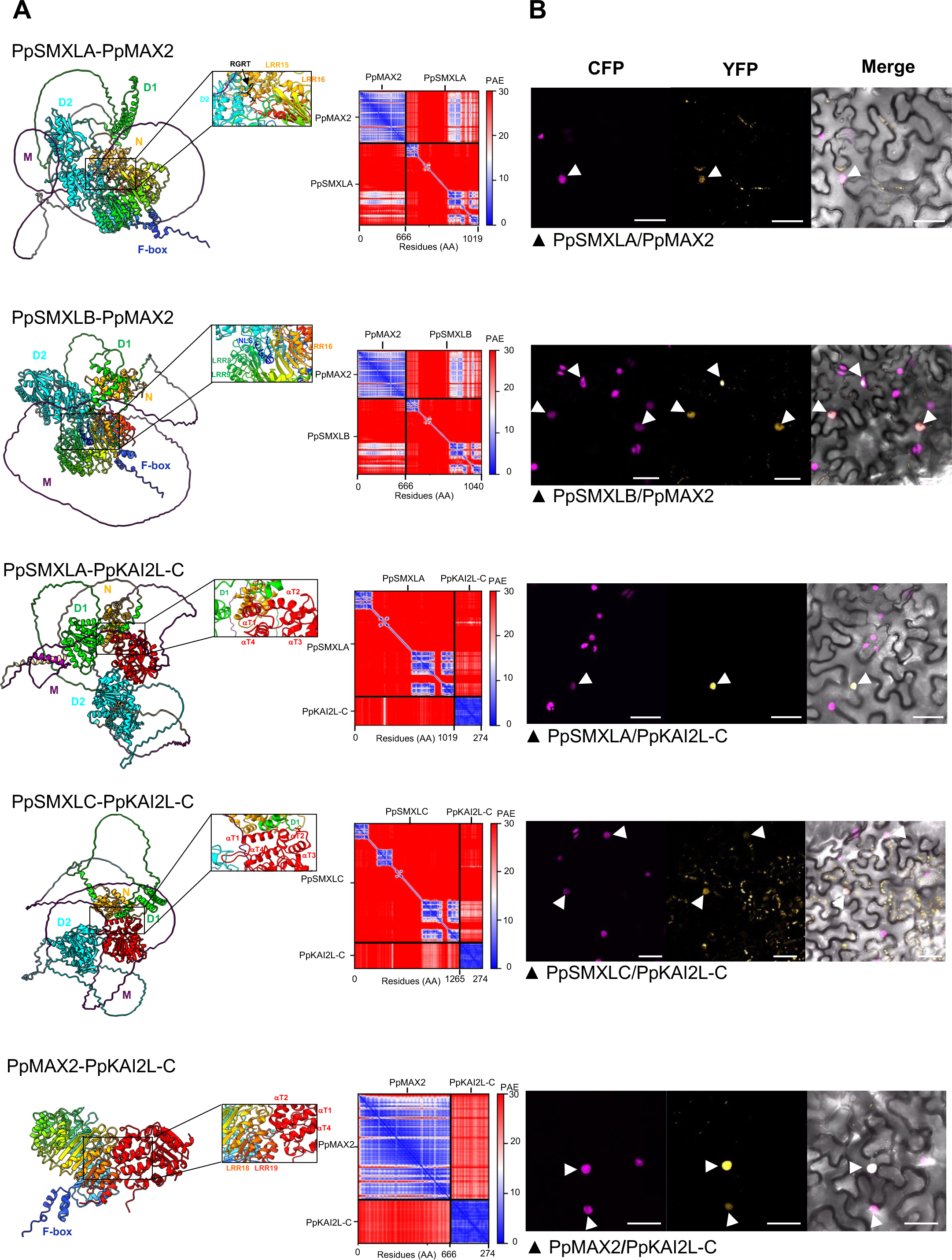
PpSMXL protein interaction predictions and assays in *Nicotiana benthamiana*. **A,** ColabFold models of, from top to bottom, PpSMXLA-PpMAX2, PpSMXLB-PpMAX2, PpSMXLA-PpKAI2L-C, PpSMXLC-PpKAI2L-C, and PpMAX2-PpKAI2L-C (AlphaFold). The PpSMXL proteins are colored by domains as used in Supplemental Figure S1, with N in orange, D1 in lime, M in purple and D2 in cyan. PpMAX2 is represented with a rainbow color palette from Nter to Cter, LRR (Leucine Rich Repeat) motifs and F-box are annotated following the rainbow palette. PpKAI2L-C is represented in red and the four alpha helices of the lid domain are annotated as αT1, αT2, αT3 and αT4 respectively. The blue dotted line symbolizes the predicted hydrogen bond between the PpSMXL RGRT motif and the residue Arg510 of the PpMAX2 LRR15 domain. Predicted Aligned Error (PAE) values for all models are shown next to the models. Low PAE values indicate strong confidence in the distances between two amino acids, and high values indicate low confidence. Insets: Interaction domains highlighted for each generated model: for PpSMXLA-PpMAX2, the RGRT (degron) motif is colored in black; for PpSMXLB-PpMAX2, a NLS (Nuclear Localization Signal) is predicted in the D2 domain and colored in dark blue. B, Bimolecular fluorescence complementation (BiFC) assays in *N. benthamiana.* PpSMXLA and PpSMXLB proteins can interact with PpMAX2, PpSMXLA and PpSMXLC proteins can interact with PpKAI2L-C, and PpMAX2 can interact with PpKAI2L-C. Below each tryptic, the first indicated protein (PpSMXLA or B) is fused to the N-terminal part of eYFP, while the second protein is fused to the C-terminal part (both tags are fused at the N-terminal end of *P. patens* proteins). Colocalization of CFP-H2b and eYFP biFC signals are pointed at with white arrowheads. Merge are overlays of CFP (magenta), YFP (yellow hot LUT) and bright field images. Bar = 50 µm. Tested interactions with DEFICIENS and GLOBOSA proteins from *Antirrhinum major* (negative controls) are shown in Supplemental Figure S16. The previously published GLOBOSA/DEFICIENS interaction (Azimzadeh et al., 2008) was consistently used a positive control of eYFP reconstruction.

Altogether, our data in *P. patens* allows us to propose a model for PpSMXL role in moss development, highlighting a novel crosstalk between the SL and KL signaling pathways (see discussion).

### Expression of PpSMXL proteins rescues SMAX4/5 function, but not SMAX1 or SMXL6/7/8 function in Arabidopsis

To investigate putative functional conservation of SMXL proteins between *Arabidopsis thaliana* and *P. patens*, we expressed PpSMXLB or PpSMXLC in Arabidopsis *smxl* mutants. In addition to the above mentioned SMAX1/SMXL2 and SMXL678 clades, Arabidopsis possesses a third subclade of SMXL proteins, SMXL345, involved in cell differentiation in a MAX2-independent manner (Wallner et al. 2017; Moturu et al. 2018; Wallner et al. 2020). We tested the functional complementation of mutants from these three clades. Expression of *PpSMXL* CDS was driven by the appropriate native Arabidopsis SMXL promoter: *pSMAX1*, *pSMXL5* and *pSMXL6*, respectively for expression in the *smax1, smxl4,5* and *smxl678* mutant. As positive controls for these complementation assays, we expressed in the Arabidopsis *smxl* mutants the native *SMAX1, SMXL5 and SMXL6* CDS under their native promoter (Supplemental Figure S17).

First, we explored the ability of PpSMXL proteins to functionally replace SMAX1 by integrating into the KL signaling pathway of Arabidopsis. Complementation was assayed by measuring hypocotyl length of seedlings grown in low light (Supplemental Figure S18, A). Only partial complementation of *smax1-2* decreased hypocotyl length was achieved by reintroducing *pSMAX1:SMAX1*, furthermore only in two out of our three independent transgenic lines (*pSMAX1:SMAX1* #20.6 and #38.1). Nonetheless, none of the plants expressing *PpSMXLB/C* showed longer hypocotyls than the untransformed *smax1-2* mutant. Interestingly, some of them developed even shorter hypocotyls (*pSMAX1:PpSMXLB* #*5.4* and *#18.3* and *pSMAX1:PpSMXLC* #*12.5* and #21.3). Thus, although the used *pSMAX1* promoter sequence may not be efficient enough, it seems reasonable to conclude that PpSMXL proteins are not able to replace SMAX1 in Arabidopsis.

Then, we explored whether PpSMXL proteins could functionally replace SMXL6 in shoot development. Plant height, caulinary secondary branch (C2) number, and leaf shape (width) are clearly affected in the *smxl678* mutant (Supplemental Figure S18, B-D). These three traits were restored to WT in the control *pSMXL6:SMXL6* lines, while none of the PpSMXLC expressing lines showed complementation. Only one line expressing PpSMXLB (#13.2) showed partial complementation of both plant height and C2 number (Supplemental Figure S18, B-C), whereas leaf width alone was partially complemented in another PpSMXLB line (#7.7). Hence, it appears PpSMXL proteins are not able to replace SMXL6 in Arabidopsis.

Finally, complementation of SMXL4/5 function was assessed by measuring the primary root length of young seedlings (Figure 7). We found that all three PpSMXLB and three out of four PpSMXLC expressing lines had similar root length as WT Arabidopsis plants and as the *smxl4,5* mutant plants complemented with native *SMXL5.* This surprising result suggests that both SMXL clades of *P. patens* have retained the same KL- and SL-independent function as Arabidopsis SMXL homologs.

**Figure 7.**
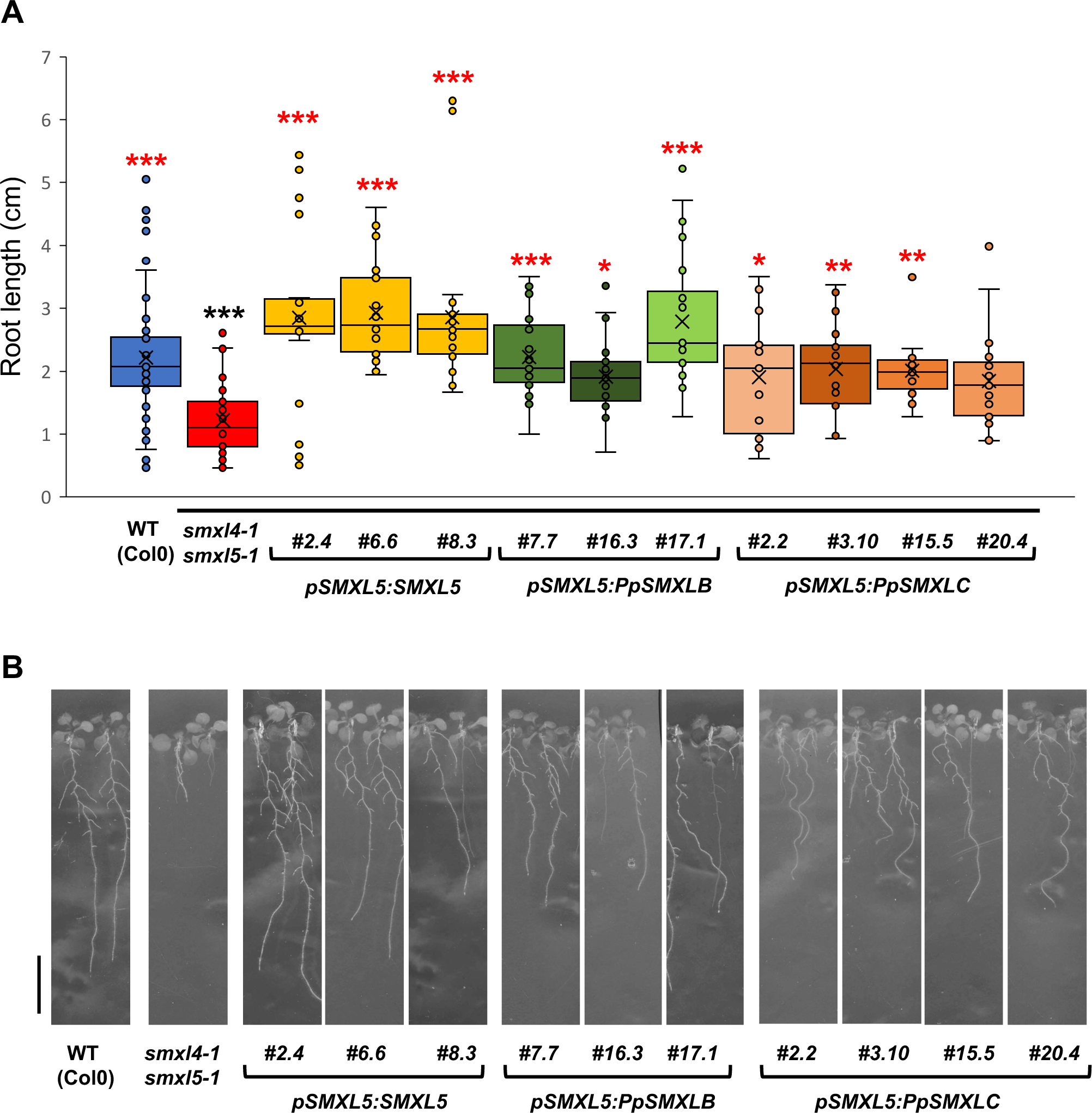
Arabidopsis *smxl4/5* mutant root phenotype is complemented by PpSMXLB and PpSMXLC. A, Root length was measured from 15 day-old seedlings grown vertically on 0.5 x MS with 1% sucrose. For control lines (Col0 and *smxl4,5*), n=38-40 seedlings per genotype, grown on 5 different plates. For all transformed lines, n=20-24 seedlings per genotype, grown on 3 different plates. Statistical significance of comparisons relative to WT is shown as black symbols and statistical significance of comparisons relative to the double mutant *smxl4,5* is shown as red symbols (Kruskal Wallis tests (both p<0.0001) followed by Dunn *post-hoc* tests). B, Representative seedlings from each genotype. Scale bar = 1 cm

## Discussion

Taken as a whole, results presented within clearly point to the PpSMXL proteins being negative regulators of growth in *P. patens*, acting downstream of PpMAX2 in the moss KL signaling pathway. Moreover, we brought some evidence that PpSMXLC/D function is required for *P. patens* to respond to exogenous SL treatment, suggesting the transduction of both KL and SL growth-regulating signals converge towards these PpSMXL proteins (dotted green line in the model in Figure 8).

**Figure 8.**
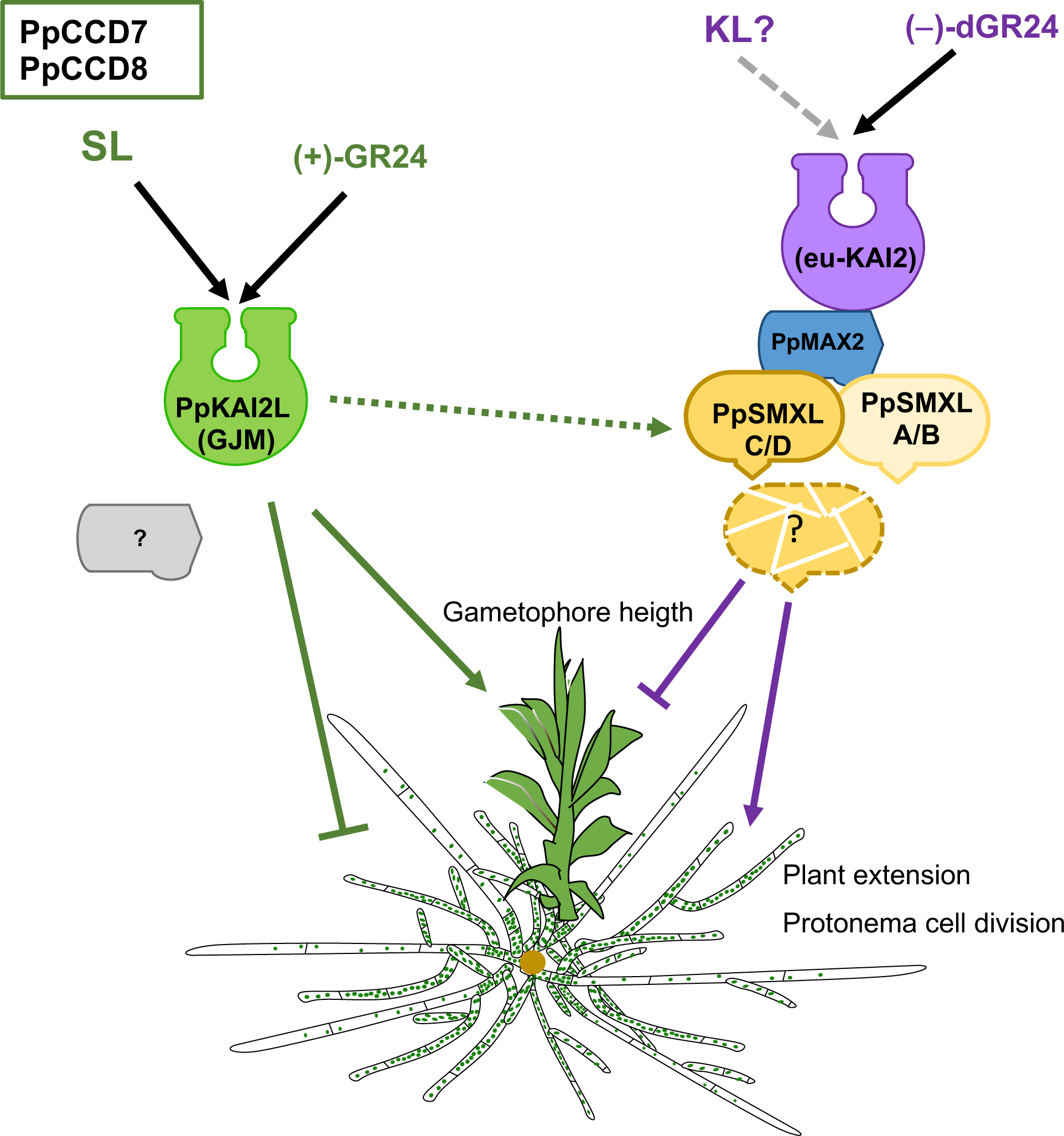
Current model of KL and SL signaling in *P. patens*. SLs and putative KL signal are mimicked by (+)-GR24 and (−)-dGR24 respectively. The two pathways have opposite effects on protonema extension and gametophore development. In contrast to angiosperms, only the KL pathway is PpMAX2 dependent. All four PpSMXL are likely suppressors of the KL pathway, though their PpMAX2-dependent degradation following KL/ (−)-dGR24 perception has not fully been demonstrated (question mark). Green dotted arrow indicates that PpSMXL could be stabilized by SLs. A further level of regulation through possible transcriptional regulation of *PpSMXL* gene transcription by PpSMXL proteins has not been represented.

### PpSMXL proteins do not act as repressors of SL response

Clade C/D *Ppsmxl* mutants do not display a constitutive SL response phenotype and instead are alike the SL deficient *Ppccd8* mutant (Figure 1; Supplemental Figure S5 and Supplemental Figure S6). This could be indicative of a positive role of PpSMXLC/D in SL signaling. Strikingly, loss-of-function of the PpSMXLA/B and/or the PpSMXLC/D clade in the *Ppccd8* background result in mutants that are like *Ppccd8* (Supplemental Figure S7). If PpSMXL proteins were repressors of SL signaling, the lift of PpSMXL-mediated repression should circumvent the absence of endogenous SL in *Ppccd8* and we could expect restoration to a WT-like phenotype. Therefore, PpSMXLs are most likely not involved in the repression of the SL response.

### PpSMXLA and PpSMXLC proteins are sensitive to a KL mimic but not to SL

We report herein that application of the KL mimic (−)-dGR24 at 10µM reduced the GFP-PpSMXLA and GFP-PpSMXLC nuclear signals in moss filaments, instead giving rise to a diffuse cytoplasmic signal (Figure 5, A-B). In addition, in a *Ppmax2-1* background, the GFP-PpSMXLA signal appeared concentrated in the nuclei. These observations lead us to hypothesize that the (−)-dGR24 effect on PpSMXL proteins may be PpMAX2-dependent.

On the other hand, the PpSMXL proteins are not rapidly degraded in response to (+)-GR24 in moss (Supplemental Figures S13B and S14). This observation is probably not the result of an insufficient amount of (+)-GR24 and/or treatment duration, as shorter treatments with similar concentrations, furthermore of the (±)-GR24 racemic mixture, have been reported to lead to complete SMXL degradation in angiosperms (Zhou et al. 2013, Wang et al. 2015, Soundappan et al. 2015). Hence, PpSMXL proteins are likely not degraded in the context of SL signaling.

Deletion of the degron motif (or degron-like for PpSMXLA/B) has little effect in *N. benthamiana* (Supplemental Figure S12), which could be explained by an incompatibility with the angiosperm MAX2 and/or with endogenous SL from *N. benthamiana*.

### PpSMXL are negative actors of the KL pathway

Four major points of evidence show that PpSMXL proteins collectively exert a negative role in the KL signaling pathway: (1) *P. patens* lines overexpressing *PpSMXLA* or *PpSMXLC* are phenotypically similar to the KL insensitive *Ppmax2-1* mutant (Figure 2); (2) Loss of function of either *PpSMXL* clade partially restores developmental disturbances caused by loss of *PpMAX2* function (Figure 3); (3) GFP-PpSMXLA and GFP-PpSMXLC fusion proteins are excluded from the nucleus and/or degraded in response to a KL mimic (Figure 5); (4) PpSMXL proteins could interact with PpMAX2 and with the putative KL receptor PpKAI2L-C (Figure 6 and Supplemental Figure S15).

As *Ppsmxl* double mutants from both clades are still able to respond to the KL mimic (−)-dGR24 (Figure 4, Supplemental Figure S9, A), the two *PpSMXL* clades likely have a redundant function in the context of KL signaling. This hypothesis, also suggested by the strong phenotype of the quadruple *smxl* mutant (Figure 1, Supplemental Figure S5), would need to be confirmed by testing the ability of higher-order mutants to respond to (−)-desmethyl-GR24, but the dramatic phenotype of these mutants makes them a challenge to work with.

### Both the KL and SL pathways regulate *PpSMXL* transcript levels

All four *PpSMXL* genes are downregulated in the *Ppmax2-1* mutant compared to WT in the dark (Supplemental Figure S10), suggesting that KL signaling would induce the transcription of these genes. This observation is comparable to what was reported for *M. polymorpha*, the unique *MpSMXL* gene being downregulated in *Mpmax2* background (Mizuno et al. 2021, and for Arabidopsis *SMAX1* and *SMXL2* genes (Stanga et al. 2013).

A negative autoregulation of *SMXL* expression was recently described for the Arabidopsis SMXL6 protein (Wang et al. 2020). A similar mechanism might exist for PpSMXL proteins, which would repress *PpSMXL* gene expression. Under our hypothesis that PpSMXL proteins are the target of PpMAX2-dependent degradation, PpSMXL protein levels are likely elevated in *Ppmax2-1*, which would lead to the observed low *PpSMXL* transcript levels in this mutant. Therefore, PpSMXL would constitute a level of negative feedback regulation in the KL pathway of *P. patens*. Moreover, *PpSMXLC/D* levels are elevated in the *Ppccd8* mutant (in light conditions, Supplemental Figure S10, C-D, middle panels) so endogenous SL response would again have an opposite effect to endogenous KL response, also at this transcriptional level. Short-term 6h treatment with the SL mimic (+)-GR24 has the expected repressive effect on *PpSMXLC/D* transcript levels, (Supplemental Figure S10, C-D, right panels).

### PpSMXL could be a bridge linking SL signaling and PpMAX2-dependent signaling

De-repression of KL signaling (*Ppsmxlcd* mutants) and restriction of SL signaling (*Ppccd8* mutant) have very similar effects on the developmental pattern of the protonema (more extended) and gametophores (smaller, especially in red light) (Figure 1). This, together with the opposite effects of (+)-GR24 (SL mimic) and (−)-desmethyl-GR24 (KL mimic) on dark-grown caulonema in WT, indicates that these two pathways could regulate the same processes in opposite manners.

In addition, we have previously shown that the *Ppmax2* mutation is epistatic to *Ppccd8* (Lopez-Obando et al. 2018). We suggested in this previous study that the phenotype of *Ppmax2* would phenocopy a constitutive response to SL. According to our present data, this phenotype would result from an over-accumulation of PpSMXL proteins. This fits our model where SL and KL signaling pathways converge on PpSMXLC/D proteins and regulate their stability/function in opposite manners (Figure 8). Indeed, as a phenotypic response to (+)-GR24 is not seen in the absence of PpSMXLC/D in some experiments, it appears that SLs act at least partially by interfering with the KL pathway. Since loss of function of either *PpMAX2* or all putative KL receptors (*PpKAI2-L A-E*) does not abolish response to (+)-GR24 (herein and (Lopez-Obando et al. 2021)), SL response does not require a functional euKAI2/PpMAX2 complex.

On the other hand, we can hypothesize that transduction of the SL signal downstream of perception by putative SL receptors (PpKAI2-L GJM) requires functional PpSMXLC/D. A tempting explanation would be that PpKAI2-L proteins of the (GJM) clade could stabilize PpSMXLC/D proteins, thus acting downstream of PpMAX2 and leading to opposite effects on phenotype compared to KL (Figure 8). We did not observe any direct interaction between PpKAI2-LG/J and PpSMXLC/D using BiFC in *N. benthamiana*. However, the hypothesis that an unknown interactor and/or presence of a *P. patens*-specific ligand is needed to permit this interaction is still open.

There might be another link between PpSMXL proteins and SL response. Indeed, we observed that triple/quadruple *Ppsmxl* mutants displayed a striking phenotype with limited protonema growth and early senescence. However, this phenotype was fully alleviated by the *Ppccd8* mutation (Supplemental Figure S7), suggesting that endogenous SLs produced via PpCCD8 activity are responsible. We previously showed that SLs have toxic effects at high concentrations on moss growth (Lopez-Obando et al. 2018). PpSMXL proteins would thus have a protective effect against SL toxicity.

### SMXL proteins likely retained a common molecular function along the evolution

We have shown through complementation assays in Arabidopsis that PpSMXLB/C expression in the *smxl4,5* mutant was enough to restore primary root length to a WT level (Figure 7). Interestingly, in Arabidopsis, SMAX1 can functionally replace SMXL5 if expressed with the same spatio-temporal profile (Wallner et al. 2017). Hence, all SMXL might have actually kept the same ancestral molecular “growth regulating” function that has been described for SMXL4/5 (Wallner et al. 2017; Wallner et al. 2020). Along evolution, this function would have (1) become KL/SL regulated for SMAX1 and SMXL678 clades through the gain of interaction with KAI2/KAI2L receptors and with MAX2, (2) become differentially regulated in various types of tissues and developmental contexts through major changes in regulatory sequences. One of the processes regulated by both SL and KL is probably cell division, as demonstrated for the *Ppccd8* mutant (Hoffmann et al. 2014). Increased cell division must be confirmed in *Ppsmxl* mutants. However, it is supported by the observation that double *Ppsmxl* mutants of both clades (tend to) develop more caulonema filaments than WT in the dark and that filaments tend to be more branched in light conditions (Figure 4; Supplemental Figure S5). *P. patens* protonema branching is inhibited by auxin (Thelander et al. 2018), but promoted by cytokinins (Cove et al. 2006). Auxin also positively regulates the chloronema to caulonema filament transition, while cytokinins positively regulate bud initiation (for a review: Guillory and Bonhomme, 2021). Four AP2-type transcriptional factors, the APB genes, are central players in this hormonal crosstalk that regulates protonema cell identity and division (Aoyama et al. 2012). Our results to date do not suggest a role for KL in chloronema to caulonema transition, but rather favor a role in bud formation (2D to 3D transition). Testing the expression of the APB genes, as well as the effect of other hormones on SL and KL related mutants, would enable to clarify their involvement at the cellular level.

The MpKAI2a/MpMAX2 pathway, regulating thallus and gemmae growth in *M. polymorpha* through the degradation of the unique MpSMXL repressor (Mizuno et al. 2021) is so far the “simplest” reported form of the above-described ancestral growth regulating function, here under the control of KL response. It is interesting to note that in *M. polymorpha*, like in *P. patens*, this pathway is regulated by light. *M. polymorpha* does not synthesize SLs, and therefore the crosstalk between SL and KL pathways that we highlight in *P. patens* cannot be observed. This crosstalk is also apparently absent in *M. paleacea* though it synthesizes BSB, likely because of the absence of the required receptor for SL as hormones (Kodama et al. 2022), though the absence of SL-sensitive SMXL copies could also play a part. Therefore, the interconnected SL and KL pathways present in *P. patens* could represent a specific case among model bryophytes species. Investigation in other bryophyte species is necessary to determine whether this SL signaling pathway is shared among mosses or a specificity of *P. patens* lineage. Moreover, in the *P. patens* model species itself, the exact nature of SL and KL signals remains to be discovered.

## Methods

### Cultivation of *Physcomitrium patens* Gransden

Unless otherwise stated in legends, experiments were always carried out on PpNO_3_ medium (minimal medium described by Ashton et al. 1979), in the following control conditions (standard growth conditions): 25°C during daytime and 23°C at night, 50% humidity, long days conditions with 16 hours of day and 8 hours of night (quantum irradiance of ∼80 µmol/m^2^/s). Multiplication of tissues from young protonema fragments prior to every experiment was done in the same conditions, using medium with higher nitrogen content (PpNH_4_ medium, PpNO_3_ medium supplemented with 2.7 mM NH_4_ tartrate). For red light experiments, plants were grown on PpNO_3_ medium in Magenta pots at 25°C, in continuous red-light (∼45 µmol µmol/m^2^/s). Cellophanes of appropriate sizes were used for monitoring of protonema extension and branching, as well as for the cultures launched in 6-well plates for gene expression studies (Guillory and Bonhomme 2021). Analysis of caulonema growth in the dark was performed in 24-well plates, with ∼2 weeks of growth in control conditions before incubation (± treatment) in the dark and placed vertically for ∼10 days (Guillory and Bonhomme 2021).

### *P. patens* gene expression analyses by qPCR

Total *P. patens* RNA was extracted and rid of contaminant genomic DNA using RNeasy Plant Mini Kit and on-column DNAse I treatment (Qiagen), following supplier’s indications. cDNA was obtained using the Maxima^TM^ H Minus retrotranscriptase (ThermoFisher), from 50-250 ng of total RNA. cDNA extracts were diluted at 1/5-1/8 before use. RT-qPCR was performed in a 384-well thermocycler (QuantStudioTM^5^, ThermoFisher), using SsoAdvanced Universal SYBR Green Supermix (BioRad) and appropriate primers. The thermocycler was programmed to run for 3 min at 95°C, followed by 40-45 cycles of 10 sec at 95°C and 30 sec at 60°C. Each biological replicate was run twice to assess technical variation. Expression of genes of interest was normalized by two reference genes among *PpElig2* (*Pp3c14_21480*), *PpAPT* (*Pp3c8_16590*) and *PpACT3* (*Pp3c10_17080*) (all three are expressed at similar levels (Le Bail et al. 2013)). Relative expression was calculated as RE = 2-CT_gene_/2-CT_ref_ where CT_ref_ is the mean value of the two reference genes. For the study of *PpSMXL* genes’ expression across development (Supplemental Figure S2B), WT *P. patens* was cultivated in Petri dishes from fragmented tissues, or in Magenta pots (for 35 days). The following tissues were collected: protonema at 6 days (mostly chloronema in our culture conditions), at 11 days (mix of chloronema and caulonema), and at 15 days (mix of chloronema, caulonema and buds), and mature gametophores and rhizoids at 35 days. Four biological replicates were used for each timepoint. For the “response to (+)-GR24” experiment (Supplemental Figure S13), plants were cultivated from fragmented *protonema* in 6-well plates for 2 weeks in control conditions, then transferred in the dark for one week, and treated with 1 µM (+)-GR24, or 0.01% DMSO in the dark for 6 hours. Six biological repeats were used for each genotype and treatment.

### Cloning of *PpSMXL* CDS

We recovered four *SMXL* genes from *Physcomitrium patens* (formerly *Physcomitrella patens*) genome on Phytozome (V3.6) as the first four results of a BLAST against *P. patens* proteome, using the full protein sequence of SMAX1 (encoded by *AT5G57710.1*). These genes were renamed *PpSMXLA* (*Pp3c2_14220*), *PpSMXLB* (*Pp3c1_23530*), *PpSMXLC* (*Pp3c9_16100*) and *PpSMXLD* (*Pp3c15_16120*). Coding sequence of each *PpSMXL* gene was amplified on WT *P. patens* Gransden cDNA, using Phusion DNA polymerase (ThermoFisher), following provided instructions and using primers with attB1 and attB2 extensions (respectively on the forward and reverse primer, see Supplemental Table S1). CDS were then integrated into the pDONR207 plasmid using BP clonase II mix (Thermofisher). pDONR207 plasmids containing *PpSMXL* CDS were submitted to PCR-mediated mutagenesis to obtain ΔRGK/RT versions.

### CRISPR-Cas9 mediated mutagenesis

Coding sequences of *PpSMXL* and *PpMAX2* were used to search for CRISPR RNA (crRNA) contiguous to a PAM motif recognized by *Streptococcus pyogenes* Cas9 (NGG), using the webtool CRISPOR V4 against *P. patens* genome Phytozome V9 (http://crispor.tefor.net/). crRNAs located in the first third of the coding sequence, with highest possible specificity score, and fewest possible predicted off-targets, were selected. Small constructs containing each crRNA fused to either the proU6 or the proU3 snRNA promoter in 5’ U3 or U6 promoter (Collonnier et al. 2017), and to the tracrRNA in 3’, encased between attB1/attB2 GateWay recombination sequences, were synthesized by Twist Biosciences. These inserts were then cloned into pDONR207 vectors. The pDONR207-sgRNA plasmids were codelivered in WT moss protoplasts with a Cas9 expressing cassette, and a cassette for transient transformation selection, through polyethylene glycol–mediated transformation, as described in (Lopez-Obando et al. 2016b). Two CRISPR-Cas9-based mutagenesis strategies were used to mutate *PpSMXL* loci in the various backgrounds (WT, *Ppccd8* or *Ppmax2-1)*. In the first strategy, we used only one guide RNA targeted against the CDS region to obtain *Ppsmxl* mutants with frameshift or small insertions/deletions (Lopez-Obando et al. 2016b). In the second strategy, crRNAs were designed in the 5’ and 3’UTR sequences, to completely remove the coding sequence of *PpSMXL* genes from the genome when used together (Supplemental Figure S1; Supplemental Table S1). Mutations giving rise to a complete deletion of the CDS were noted as *Ppsmxl*Δ (Supplemental Figure S3 and S4). Mutations obtained from the first strategy were genotyped by PCR amplification of *PpSMXL* loci around the recognition sequence of each guide RNA and sequencing of the PCR products. Those obtained from the second strategy were genotyped by monitoring the size and sequence of amplicons spanning from the 5’UTR to the 3’UTR. *Ppsmxlab* mutants in the *Ppmax2-1* background were obtained, but no *Ppsmxlcd* mutants. Therefore, we instead mutated *PpMAX2* in one of the *Ppsmxl*Δ*cd* mutants, using the CRISPR-Cas9 system, with five guide RNAs targeting *PpMAX2* (Supplemental Figure S10, Supplemental Table S1). Using this approach, we successfully obtained a new *Ppmax2* mutant allele (*Ppmax2-16*) in the *Ppsmxl*Δ*cd* mutant background.

### Generation of proZmUbi:GFP-PpSMXL and control proZmUbi:flag-GFP lines

*proZmUbi:GFP-PpSMXL* constructs were obtained by LR recombination of pDONR207 plasmids containing *PpSMXL* coding sequences with the pMP1335 destination vector (http://labs.biology.ucsd.edu/estelle/Moss_files/pK108N+Ubi-mGFP6-GW.gb). These plasmids were used independently to transform WT *P. patens* Gransden, together with pDONR207 containing sgRNA recognizing Pp108 homology sequences contained in the three pMP vectors and appropriate Cas9 and selection plasmids (Lopez-Obando et al. 2016b). Obtained G418 resistant lines were screened for insertion using PCR (with proPpSMXL forward and GUS reverse, GFP forward and PpSMXL reverse, or proZmUbi forward and GFP reverse primers, respectively, Supplemental Table S1).

### Generation of BiFC constructs

Gateway cloned inserts of genes of interest were integrated into pbiFP vectors using LR Clonase II mix (ThermoFisher). Inserts containing a STOP codon were cloned in pbiFP2 and pbiFP3, those not containing a STOP were cloned in pbiFP1 and pbiFP4 (when possible, all four vectors were obtained for a given gene). Resulting vectors were electroporated into *Escherichia coli* DH10B cells and clones were selected on spectinomycin. In-frame integration of the coding sequence relative to the half eYFP tag was checked by sequencing insert’s ends. The previously published GLOBOSA/DEFICIENS interaction (Azimzadeh et al. 2008) was consistently used as a positive control of eYFP reconstruction.

### Agroinfiltration of *Nicotiana benthamiana* leaves

pbiFP plasmids containing the genes of interest were electroporated into *Agrobacterium tumefaciens* strain C58C1. Agrobacteria were incubated for 18 hours at 28°C under constant agitation and then pelleted, washed twice, and resuspended in infiltration buffer (13 g/L S-medium (Duchefa Biochemie) and 40 g/L sucrose, pH 5.7 with KOH) to attain an OD600 value of 0.5. To enhance transient p35S-driven expression of RFP-PpSMXL and BiFC fusion proteins, the P19 viral suppressor of gene silencing from tomato bushy stunt virus was co-expressed. Equal volumes of needed bacterial cultures were mixed and infiltrated into the abaxial epidermis of 4–5-week-old *Nicotiana benthamiana* leaves (in a line stably expressing CFP tagged H2b histone). After incubation at 25 °C (16 h light/8 h dark) for 4 days, leaves were harvested on wet paper and kept in similar temperature and hygrometry conditions for short-term preservation until observation.

### Confocal microscopy observations

Fragments of *P. patens proZmUBI:GFP-PpSMXL* plants and infiltrated parts of *Nicotiana benthamiana* leaves were both observed on a TCS SP5 inverted or on a TCS SP8 Upright confocal microscopy system (Leica), with a 20X objective. GFP fluorescence was acquired in the 495nm-520nm λ range, eYFP in the 525nm-540nm range, RFP in the 570nm-610nm range and CFP in the 465nm-505nm range. Signals in the 700nm-750nm range were attributed to chlorophyll autofluorescence. Lasers used for excitation have a peak wavelength of 488nm (GFP), 514nm (YFP), 458nm (CFP) and 561nm (RFP).

### AlphaFold2 and ColabFold predictions

AlphaFold2 predictions of both monomers and multimers were computed through the ColabFold notebook (ColabFold v1.3 and AlphaFold2 v2.2) using a ColabPro+ plan (https://colab.research.google.com/github/sokrypton/ColabFold/blob/main/AlphaFold2.ipynb). The plDDT, PAE scores and graphs were provided directly by the provided online notebook (Mirdita et al. 2022). Predicted structures were not relaxed using amber, and no template information was used. mmseqs_uniref_env was used for the unpaired MSA, and sequences from the same species were paired. In advanced settings, automatic modes were kept. Images were produced using UCSF ChimeraX (Pettersen et al. 2021).

### Constructs and generation of Arabidopsis transgenic lines

The expression vectors for transgenic Arabidopsis were constructed by MultiSite Gateway Three-Fragment Vector Construction kit (Invitrogen). SMAX/SMXL and PpSMXL constructs were tagged with m-Citrine protein at their C-terminus. Lines were resistant to hygromycin. For the Arabidopsis *SMAX1*, *SMXL5*, and *SMXL6* promotors cloning, a 3128-bp fragment upstream from SMAX1 start codon, a 3028-bp fragment upstream from SMXL5 start codon and a 3058-bp fragment upstream from SMXL6 start codon were amplified from Col-0 genomic DNA using the primers described in Supplemental Table S1, and were cloned into the pDONR-P4P1R vector, using Gateway recombination (Invitrogen) as described in (Lopez-Obando et al. 2021). The m-Citrine tag was cloned into pDONR-P2RP3 (Invitrogen) as described in (de Saint Germain et al. 2016). SMAX1, AtSMXL5 and AtSMXL6 CDS were PCR amplified from Arabidopsis Col-0 cDNA and with the primers specified in Supplemental Table S1 and then recombined into the pDONR221 vector (Invitrogen). PpSMXLB and PpSMXLC CDS were obtained as described above. The suitable combinations of AtSMXL native promoters, AtSMXL or PpSMXL CDS, and m-Citrine tag were cloned into the pH7m34GW final destination vectors by using the three fragment recombination system (Karimi et al. 2007) and were thusly named *pSMAX1:SMAX1, pSMAX1:PpSMXLB, pSMAX1:PpSMXLC, pSMXL5:SMXL5, pSMXL5:PpSMXLB, pSMXL5:PpSMXLC, pSMXL6:SMXL6, pSMXL6:PpSMXLB and pSMXL6:PpSMXLC*. Transformation of Arabidopsis *smax1-2*, *smxl4-1,5-1* and *smxl678* mutants was performed according to the conventional floral dipping method (Clough and Bent 1998), with Agrobacterium strain GV3101. For each construct, at least 12 independent T1 lines were isolated, and then 2 to 4 lines were selected in T2 for showing a 3 :1 segregation (single T-DNA insertion). Phenotypic analysis shown in Figure 8 and Supplemental Figures S20 were performed on the T3 homozygous lines.

### Arabidopsis plant materials, growth conditions, and phenotypic assays

Arabidopsis plants used in this study originated from the Columbia (Col-0) ecotype background. *smax1-2* and *smxl678* Arabidopsis mutants are gift from Dave Nelson (University of California, Riverside), and the Arabidopsis *smxl4-1 smxl5-1* double mutant gift from Thomas Greb (Heidelberg University, Heidelberg). For shoot branching assays, the plants were grown in greenhouse. Experiments were carried out in summer, under long photoperiods (15-16 h per day); daily temperatures fluctuated between 18 °C and 25 °C. Peak levels of PAR were between 700 and 1000 μmol m^-2^ s^-1^. Plants were watered twice a week with tap water. The number of caulinary branch and plant height were scored when the plants were 35 days old. Leaf Morphology Assay was performed as described in (Soundappan et al. 2015) on the 5th leaf of each plant marked with indelible marker at 3 weeks post-germination. Plants were cultivated as for branching assay. Hypocotyl elongation assays were performed as described in (Guercio et al. 2022) with the following modification: 11-day-old seedlings were photographed, and hypocotyl lengths were quantified using ImageJ. 2 plates of 20 to 24 seeds were sown for each genotype. For root length assay, seedlings were grown on 3 different plates in the same media and condition as for hypocotyl assay. Primary root length was measured at 15 days post-germination using ImageJ.

### Chemicals

GR24 enantiomers and (−)-desmethyl-GR24 were produced by V Steinmetz and F-D Boyer using organic synthesis and chiral separation as described in (de Saint Germain et al. 2021). Chemicals were diluted in DMSO.

### RT-PCR analysis in Arabidopsis

Semi-quantitative RT-PCR analyses of Arabidopsis transformants were performed from leaf or seedling RNAs, extracted and rid of contaminant genomic DNA using RNeasy Plant Mini Kit and on-column DNAse I treatment (Qiagen), following supplier’s indications. cDNA was obtained using the H Minus retrotranscriptase (ThermoFisher), from 250 ng of total RNA. cDNA extracts were diluted twice, and 1 μL was used for each PCR reaction. The thermocycler was programmed to run for 3 min at 95 °C, followed by 28 cycles of 30 sec at 95 °C, 30 sec at 56 °C and 40 sec at 72 °C.

### Statistical analysis of results

Kruskal-Wallis, Mann-Whitney and *post-hoc* Dunn, Dunnett or Tukey multiple comparisons tests (details in figures legends) were carried out either in R 3.6.3 or in GraphPad Prism 8.4.2. Tests employed were mostly non-parametric as normality of distributions and/or homoscedasticity among groups could not be confirmed in most experiments (Kruskal-Wallis tests for multiple comparisons and Mann-Whitney for single comparisons, unless otherwise stated in legends). For some gene expression experiments, data points were excluded based on an outliers’ search (Grubb’s, α=0.05) on in GraphPad Prism 8.4.2. Unless otherwise defined, used statistical significance scores are as follow: # 0.05≤p<0.1, * 0.01≤p<0.05, ** 0.001≤p<0.01, *** p<0.001. Same letters scores indicate that p≥0.05 (non-significant differences).

### Accession Numbers

Sequences used in the present article can be found on Phytozome (*P. patens* Gransden genome, V3.1 version). *PpSMXLA* is *Pp3c2_14220, PpSMXLB* is *Pp3c1_23530*, *PpSMXLC* is *Pp3c9_16100* and *PpSMXLD* is *Pp3c15_16120. PpMAX2* corresponds to *Pp3c17_1180*, *PpCCD8 to Pp3c6_21520*, *PpAPT* to *Pp3c8_16590, PpACT3* to *Pp3c10_17080,* and *PpElig2* corresponds to *Pp3c14_21480*.

### Authors contributions

A.G performed most of the experiments. Some steps in genotyping mutant and transgenic lines and support with molecular cloning was given by P.L.B. K.B, A.L and S.B helped for moss mutant phenotyping and J-P. P for transgenic Arabidopsis phenotyping. V.S and F-D. B furnished SL and KL analogs. Plasmids used for *Ppsmxl* mutant generation in the first strategy, as well as higher order *Ppsmxl* mutants were obtained by M.L-O. L.L generated AlphaFold2-based models for protein interactions. A.G, S.B, A.S.G, and C.R all contributed to experimental design and to the argumentation developed herein.

## Acknowledgements

We thank Dave Nelson (University of California, Riverside) for the generous gift of Arabidopsis *smax1-2* and *smxl678* mutants, as well as for the pSMXL6:SMXL6-GFP construct. We thank Thomas Greb (Heidelberg University, Heidelberg) for the kind gift of Arabidopsis *smxl4,5* double mutant. We thank Fabien Nogué (IJPB, Versailles) for his helpful advice and for giving us guide RNAs targeting the Pp108 non-coding locus. We thank Bastien Cayrol for his participation in several assays during his Master 1 internship. We thank Martine Pastuglia (IJPB, Versailles) for the gift of BiFC control constructs and Michael J. Prigge (University of California San Diego, La Jolla) for nicely sending us the pMP destination vectors. This work has benefited from the support of IJPB’s Plant Observatory technological platforms.

## Fundings

The IJPB benefits from the support of Saclay Plant Sciences-SPS (ANR-17-EUR-0007). This work has benefited from a French State grant (Saclay Plant Sciences, reference n° ANR-17-EUR-0007, EUR SPS-GSR) managed by the French National Research Agency under an Investments for the Future program integrated into France 2030 (reference n° ANR-11-IDEX-0003-02).

## Notes

### Competing Interest Statement

The authors have declared no competing interest.

### Summary of Updates

The text and several figures have been revised according to reviewiers comments.

## References

Akiyama K, Matsuzaki K, Hayashi H (2005) Plant sesquiterpenes induce hyphal branching in arbuscular mycorrhizal fungi. Nature 435 (7043):824–827. doi:10.1038/nature03608

Alder A, Jamil M, Marzorati M, Bruno M, Vermathen M, Bigler P, Ghisla S, Bouwmeester H, Beyer P, Al-Babili S (2012) The path from beta-carotene to carlactone, a strigolactone-like plant hormone. Science 335 (6074):1348–1351. doi:10.1126/science.1218094

Aoyama T, Hiwatashi Y, Shigyo M, Kofuji R, Kubo M, Ito M, Hasebe M (2012) AP2-type transcription factors determine stem cell identity in the moss *Physcomitrella patens*. Development 139(17):3120–3129. doi: 10.1242/dev.076091

Azimzadeh J, Nacry P, Christodoulidou A, Drevensek S, Camilleri C, Amiour N, Parcy F, Pastuglia M, Bouchez D (2008) Arabidopsis TONNEAU1 proteins are essential for preprophase band formation and interact with centrin. Plant Cell 20 (8):2146–2159. doi:tpc.107.056812 [pii]10.1105/tpc.107.056812

Bennett T, Liang Y, Seale M, Ward S, Muller D, Leyser O (2016) Strigolactone regulates shoot development through a core signalling pathway. Biol Open 5 (12):1806–1820. doi:10.1242/bio.021402

Besserer A, Becard G, Jauneau A, Roux C, Sejalon-Delmas N (2008) GR24, a synthetic analog of strigolactones, stimulates the mitosis and growth of the arbuscular mycorrhizal fungus Gigaspora rosea by boosting its energy metabolism. Plant Physiol 148 (1):402–413

Besserer A, Puech-Pages V, Kiefer P, Gomez-Roldan V, Jauneau A, Roy S, Portais JC, Roux C, Becard G, Sejalon-Delmas N (2006) Strigolactones stimulate arbuscular mycorrhizal fungi by activating mitochondria. PLoS Biol 4 (7):e226

Bianchi A, Giorgi C, Ruzza P, Toniolo C, Milner-White EJ (2012) A synthetic hexapeptide designed to resemble a proteinaceous P-loop nest is shown to bind inorganic phosphate. Proteins 80 (5):1418–1424. doi:10.1002/prot.24038

Bonhomme S, Guillory A (2022) Synthesis and Signalling of Strigolactone and KAI2-Ligand signals in bryophytes. J Exp Bot. doi:10.1093/jxb/erac186

Bythell-Douglas R, Rothfels CJ, Stevenson DWD, Graham SW, Wong GK, Nelson DC, Bennett T (2017) Evolution of strigolactone receptors by gradual neo-functionalization of KAI2 paralogues. BMC biology 15 (1):52. doi:10.1186/s12915-017-0397-z

Choi J, Lee T, Cho J, Servante EK, Pucker B, Summers W, Bowden S, Rahimi M, An K, An G, Bouwmeester HJ, Wallington EJ, Oldroyd G, Paszkowski U (2020) The negative regulator SMAX1 controls mycorrhizal symbiosis and strigolactone biosynthesis in rice. Nat Commun 11 (1):2114. doi:10.1038/s41467-020-16021-1

Clough SJ, Bent AF (1998) Floral dip: a simplified method for Agrobacterium-mediated transformation of Arabidopsis thaliana. Plant J 16 (6):735–743. doi:10.1046/j.1365-313x.1998.00343.x

Collonnier C, Guyon-Debast A, Maclot F, Mara K, Charlot F, Nogué F (2017) Towards mastering CRISPR-induced gene knock-in in plants: Survey of key features and focus on the model Physcomitrella patens. Methods 121–122:103-117. doi:10.1016/j.ymeth.2017.04.024

Conn CE, Nelson DC (2016) Evidence that KARRIKIN-INSENSITIVE2 (KAI2) Receptors may Perceive an Unknown Signal that is not Karrikin or Strigolactone. Front Plant Sci 6:1219. doi:10.3389/fpls.2015.01219

Cook CE, Whichard LP, Turner B, Wall ME, Egley GH (1966) Germination of Witchweed (Striga lutea Lour.): Isolation and Properties of a Potent Stimulant. Science 154 (3753):1189–1190

Coudert Y, Palubicki W, Ljung K, Novak O, Leyser O, Harrison CJ (2015) Three ancient hormonal cues co-ordinate shoot branching in a moss. Elife 4. doi:10.7554/eLife.06808

Cove D, Bezanilla M, Harries P, Quatrano R (2006) Mosses as model systems for the study of metabolism and development. Annu Rev Plant Biol. 57:497–520. doi: 10.1146/annurev.arplant.57.032905.105338

de Saint Germain A, Clave G, Badet-Denisot MA, Pillot JP, Cornu D, Le Caer JP, Burger M, Pelissier F, Retailleau P, Turnbull C, Bonhomme S, Chory J, Rameau C, Boyer FD (2016) An histidine covalent receptor and butenolide complex mediates strigolactone perception. Nat Chem Biol 12 (10):787–794. doi:10.1038/nchembio.2147

de Saint Germain A, Jacobs A, Brun G, Pouvreau JB, Braem L, Cornu D, Clavé G, Baudu E, Steinmetz V, Servajean V, Wicke S, Gevaert K, Simier P, Goormachtig S, Delavault P, Boyer FD (2021) A Phelipanche ramosa KAI2 protein perceives strigolactones and isothiocyanates enzymatically. Plant Commun 2 (5):100166. doi:10.1016/j.xplc.2021.100166

Decker EL, Alder A, Hunn S, Ferguson J, Lehtonen MT, Scheler B, Kerres KL, Wiedemann G, Safavi-Rizi V, Nordzieke S, Balakrishna A, Baz L, Avalos J, Valkonen JPT, Reski R (2017) Strigolactone biosynthesis is evolutionarily conserved, regulated by phosphate starvation and contributes to resistance against phytopathogenic fungi in a moss, Physcomitrella patens. New Phytol 216 (2):455–468. doi:10.1111/nph.14506

Delaux PM, Xie X, Timme RE, Puech-Pages V, Dunand C, Lecompte E, Delwiche CF, Yoneyama K, Becard G, Sejalon-Delmas N (2012) Origin of strigolactones in the green lineage. New Phytol 195 (4):857–871. doi:10.1111/j.1469-8137.2012.04209.x

Gomez-Roldan V, Fermas S, Brewer PB, Puech-Pages V, Dun EA, Pillot JP, Letisse F, Matusova R, Danoun S, Portais JC, Bouwmeester H, Becard G, Beveridge CA, Rameau C, Rochange SF (2008) Strigolactone inhibition of shoot branching. Nature 455 (7210):189–194. doi:10.1038/nature07271

Guercio AM, Torabi S, Cornu D, Dalmais M, Bendahmane A, Le Signor C, Pillot JP, Le Bris P, Boyer FD, Rameau C, Gutjahr C, de Saint Germain A, Shabek N (2022) Structural and functional analyses explain Pea KAI2 receptor diversity and reveal stereoselective catalysis during signal perception. Commun Biol 5 (1):126. doi:10.1038/s42003-022-03085-6

Guillory A, Bonhomme S (2021) Methods for Medium-Scale Study of Biological Effects of Strigolactone-Like Molecules on the Moss Physcomitrium (Physcomitrella) patens. In: Prandi C, Cardinale F (eds) Methods Mol Biol, vol 2309. 2021/05/25 edn. Springer US, pp 143–155. doi:10.1007/978-1-0716-1429-7_12

Hoffmann B, Proust H, Belcram K, Labrune C, Boyer FD, Rameau C, Bonhomme S (2014) Strigolactones inhibit caulonema elongation and cell division in the moss Physcomitrella patens. PLoS One 9 (6):e99206. doi:10.1371/journal.pone.0099206

Jiang L, Liu X, Xiong G, Liu H, Chen F, Wang L, Meng X, Liu G, Yu H, Yuan Y, Yi W, Zhao L, Ma H, He Y, Wu Z, Melcher K, Qian Q, Xu HE, Wang Y, Li J (2013) DWARF 53 acts as a repressor of strigolactone signalling in rice. Nature 504 (7480):401–405. doi:10.1038/nature12870

Jones DT, Taylor WR, Thornton JM (1992). The rapid generation of mutation data matrices from protein sequences. Computer Applications in the Biosciences 8: 275–282.

Jumper J, Evans R, Pritzel A, Green T, Figurnov M, Ronneberger O, Tunyasuvunakool K, Bates R, Žídek A, Potapenko A, Bridgland A, Meyer C, Kohl SAA, Ballard AJ, Cowie A, Romera-Paredes B, Nikolov S, Jain R, Adler J, Back T, Petersen S, Reiman D, Clancy E, Zielinski M, Steinegger M, Pacholska M, Berghammer T, Bodenstein S, Silver D, Vinyals O, Senior AW, Kavukcuoglu K, Kohli P, Hassabis D (2021). Highly accurate protein structure prediction with AlphaFold. Nature 596:583–589. 10.1038/s41586-021-03819-2.

Karimi M, Bleys A, Vanderhaeghen R, Hilson P (2007) Building blocks for plant gene assembly. Plant Physiol 145 (4):1183–1191. doi:10.1104/pp.107.110411

Kerr SC, Patil SB, de Saint Germain A, Pillot JP, Saffar J, Ligerot Y, Aubert G, Citerne S, Bellec Y, Dun EA, Beveridge CA, Rameau C (2021) Integration of the SMXL/D53 strigolactone signalling repressors in the model of shoot branching regulation in Pisum sativum. Plant J 107 (6):1756–1770. doi:10.1111/tpj.15415

Khosla A, Morffy N, Li Q, Faure L, Chang SH, Yao J, Zheng J, Cai ML, Stanga J, Flematti GR, Waters MT, Nelson DC (2020) Structure-Function Analysis of SMAX1 Reveals Domains That Mediate Its Karrikin-Induced Proteolysis and Interaction with the Receptor KAI2. Plant Cell 32 (8):2639–2659. doi:10.1105/tpc.19.00752

Kodama K, Rich MK, Yoda A, Shimazaki S, Xie X, Akiyama K, Mizuno Y, Komatsu A, Luo Y, Suzuki H, Kameoka H, Libourel C, Keller J, Sakakibara K, Nishiyama T, Nakagawa T, Mashiguchi K, Uchida K, Yoneyama K, Tanaka Y, Yamaguchi S, Shimamura M, Delaux PM, Nomura T, Kyozuka J (2022) An ancestral function of strigolactones as symbiotic rhizosphere signals. Nat Commun 13 (1):3974. doi:10.1038/s41467-022-31708-3

Kyozuka J, Nomura T, Shimamura M (2022) Origins and evolution of the dual functions of strigolactones as rhizosphere signaling molecules and plant hormones. Curr Opin Plant Biol 65:102154. doi:10.1016/j.pbi.2021.102154

Kumar S., Stecher G., Li M., Knyaz C., and Tamura K. (2018). MEGA X: Molecular Evolutionary Genetics Analysis across computing platforms. Molecular Biology and Evolution 35:1547–1549.

Le Bail A, Scholz S, Kost B (2013) Evaluation of reference genes for RT qPCR analyses of structure-specific and hormone regulated gene expression in Physcomitrella patens gametophytes. PLoS One 8 (8):e70998. doi:10.1371/journal.pone.0070998

Liang Y, Ward S, Li P, Bennett T, Leyser O (2016) SMAX1-LIKE7 Signals from the Nucleus to Regulate Shoot Development in Arabidopsis via Partially EAR Motif-Independent Mechanisms. Plant Cell 28 (7):1581–1601. doi:10.1105/tpc.16.00286

Lopez-Obando M, Conn CE, Hoffmann B, Bythell-Douglas R, Nelson DC, Rameau C, Bonhomme S (2016a) Structural modelling and transcriptional responses highlight a clade of PpKAI2-LIKE genes as candidate receptors for strigolactones in Physcomitrella patens. Planta 243 (6):1441–1453. doi:10.1007/s00425-016-2481-y

Lopez-Obando M, de Villiers R, Hoffmann B, Ma L, de Saint Germain A, Kossmann J, Coudert Y, Harrison CJ, Rameau C, Hills P, Bonhomme S (2018) Physcomitrella patens MAX2 characterization suggests an ancient role for this F-box protein in photomorphogenesis rather than strigolactone signalling. New Phytol 219 (2):743–756. doi:10.1111/nph.15214

Lopez-Obando M, Guillory A, Boyer FD, Cornu D, Hoffmann B, Le Bris P, Pouvreau JB, Delavault P, Rameau C, de Saint Germain A, Bonhomme S (2021) The Physcomitrium (Physcomitrella) patens PpKAI2L receptors for strigolactones and related compounds function via MAX2-dependent and -independent pathways. Plant Cell 33 (11):3487–3512. doi:10.1093/plcell/koab217

Lopez-Obando M, Hoffmann B, Géry C, Guyon-Debast A, Téoulé E, Rameau C, Bonhomme S, Nogué F (2016b) Simple and Efficient Targeting of Multiple Genes Through CRISPR-Cas9 in Physcomitrella patens. G3 (Bethesda) 6 (11):3647-3653. doi:10.1534/g3.116.033266

Ma H, Duan J, Ke J, He Y, Gu X, Xu TH, Yu H, Wang Y, Brunzelle JS, Jiang Y, Rothbart SB, Xu HE, Li J, Melcher K (2017) A D53 repression motif induces oligomerization of TOPLESS corepressors and promotes assembly of a corepressor-nucleosome complex. Science advances 3 (6):e1601217. doi:10.1126/sciadv.1601217

Machin DC, Hamon-Josse M, Bennett T (2020) Fellowship of the rings: a saga of strigolactones and other small signals. New Phytol 225 (2):621–636. doi:10.1111/nph.16135

Mirdita M, Schütze K, Moriwaki Y, Heo L, Ovchinnikov S, Steinegger M (2022) ColabFold: making protein folding accessible to all. Nat Methods 19: 679–682 doi:10.1038/s41592-022-01488-1

Mizuno Y, Komatsu A, Shimazaki S, Naramoto S, Inoue K, Xie X, Ishizaki K, Kohchi T, Kyozuka J (2021) Major components of the KARRIKIN INSENSITIVE2-dependent signaling pathway are conserved in the liverwort Marchantia polymorpha. Plant Cell 33 (7):2395–2411. doi:10.1093/plcell/koab106

Moturu TR, Thula S, Singh RK, Nodzynski T, Vareková RS, Friml J, Simon S (2018) Molecular evolution and diversification of the SMXL gene family. J Exp Bot 69 (9):2367–2378. doi:10.1093/jxb/ery097

Ortiz-Ramirez C, Hernandez-Coronado M, Thamm A, Catarino B, Wang M, Dolan L, Feijo JA, Becker JD (2016) A Transcriptome Atlas of Physcomitrella patens Provides Insights into the Evolution and Development of Land Plants. Mol Plant 9 (2):205–220. doi:10.1016/j.molp.2015.12.002

Park YJ, Kim JY, Park CM (2022) SMAX1 potentiates phytochrome B-mediated hypocotyl thermomorphogenesis. Plant Cell 34 (7):2671–2687. doi:10.1093/plcell/koac124

Pettersen EF, Goddard TD, Huang CC, Meng EC, Couch GS, Croll TI, Morris JH, Ferrin TE (2021) UCSF ChimeraX: Structure visualization for researchers, educators, and developers. Protein Sci. (1):70-82. doi: 10.1002/pro.3943.

Proust H, Hoffmann B, Xie X, Yoneyama K, Schaefer DG, Nogue F, Rameau C (2011) Strigolactones regulate protonema branching and act as a quorum sensing-like signal in the moss Physcomitrella patens. Development 138 (8):1531–1539. doi:10.1242/dev.058495

Radhakrishnan GV, Keller J, Rich MK, Vernié T, Mbadinga Mbadinga DL, Vigneron N, Cottret L, Clemente HS, Libourel C, Cheema J, Linde AM, Eklund DM, Cheng S, Wong GKS, Lagercrantz U, Li FW, Oldroyd GED, Delaux PM (2020) An ancestral signalling pathway is conserved in intracellular symbioses-forming plant lineages. Nat Plants 6 (3):280–289. doi:10.1038/s41477-020-0613-7

Soundappan I, Bennett T, Morffy N, Liang Y, Stanga JP, Abbas A, Leyser O, Nelson D (2015) SMAX1-LIKE/D53 Family Members Enable Distinct MAX2-Dependent Responses to Strigolactones and Karrikins in Arabidopsis. Plant Cell 27 (11):3143–3159. doi:10.1105/tpc.15.00562

Stanga JP, Morffy N, Nelson DC (2016) Functional redundancy in the control of seedling growth by the karrikin signaling pathway. Planta 243 (6):1397–1406. doi:10.1007/s00425-015-2458-2

Stanga JP, Smith SM, Briggs WR, Nelson DC (2013) SUPPRESSOR OF MORE AXILLARY GROWTH2 1 controls seed germination and seedling development in Arabidopsis. Plant Physiol 163 (1):318–330. doi:10.1104/pp.113.221259

Sun YK, Flematti GR, Smith SM, Waters MT (2016) Reporter Gene-Facilitated Detection of Compounds in Arabidopsis Leaf Extracts that Activate the Karrikin Signaling Pathway. Front Plant Sci 7:1799. doi:10.3389/fpls.2016.01799

Temmerman A, Guillory A, Bonhomme S, Goormachtig S, Struk S (2022) Masks Start to Drop: Suppressor of MAX2 1-Like Proteins Reveal Their Many Faces. Front Plant Sci 13:887232. doi:10.3389/fpls.2022.887232

Thelander M, Landberg K, Sundberg E (2018) Auxin-mediated developmental control in the moss *Physcomitrella patens*. J Exp Bot. 69(2):277–290. doi: 10.1093/jxb/erx255

Umehara M, Hanada A, Yoshida S, Akiyama K, Arite T, Takeda-Kamiya N, Magome H, Kamiya Y, Shirasu K, Yoneyama K, Kyozuka J, Yamaguchi S (2008) Inhibition of shoot branching by new terpenoid plant hormones. Nature 455 (7210):195–200. doi:10.1038/nature07272

Villaecija-Aguilar JA, Hamon-Josse M, Carbonnel S, Kretschmar A, Schmidt C, Dawid C, Bennett T, Gutjahr C (2019) SMAX1/SMXL2 regulate root and root hair development downstream of KAI2-mediated signalling in Arabidopsis. PLoS genetics 15 (8):e1008327. doi:10.1371/journal.pgen.1008327

Walker CH, Siu-Ting K, Taylor A, O’Connell MJ, Bennett T (2019) Strigolactone synthesis is ancestral in land plants, but canonical strigolactone signalling is a flowering plant innovation. BMC biology 17 (1):70. doi:10.1186/s12915-019-0689-6

Wallner ES, López-Salmerón V, Belevich I, Poschet G, Jung I, Grünwald K, Sevilem I, Jokitalo E, Hell R, Helariutta Y, Agustí J, Lebovka I, Greb T (2017) Strigolactone- and Karrikin-Independent SMXL Proteins Are Central Regulators of Phloem Formation. Curr Biol 27 (8):1241–1247. doi:10.1016/j.cub.2017.03.014

Wallner ES, Tonn N, Shi D, Jouannet V, Greb T (2020) SUPPRESSOR OF MAX2 1-LIKE 5 promotes secondary phloem formation during radial stem growth. Plant J 102 (5):903–915. doi:10.1111/tpj.14670

Wang L, Wang B, Jiang L, Liu X, Li X, Lu Z, Meng X, Wang Y, Smith SM, Li J (2015) Strigolactone Signaling in Arabidopsis Regulates Shoot Development by Targeting D53-Like SMXL Repressor Proteins for Ubiquitination and Degradation. Plant Cell 27 (11):3128–3142. doi:10.1105/tpc.15.00605

Wang L, Wang B, Yu H, Guo H, Lin T, Kou L, Wang A, Shao N, Ma H, Xiong G, Li X, Yang J, Chu J, Li J (2020) Transcriptional regulation of strigolactone signalling in Arabidopsis. Nature 583 (7815):277–281. doi:10.1038/s41586-020-2382-x

Waters MT, Scaffidi A, Moulin SL, Sun YK, Flematti GR, Smith SM (2015) A Selaginella moellendorffii Ortholog of KARRIKIN INSENSITIVE2 Functions in Arabidopsis Development but Cannot Mediate Responses to Karrikins or Strigolactones. Plant Cell 27 (7):1925–1944. doi:10.1105/tpc.15.00146

Yao J, Scaffidi A, Meng Y, Melville KT, Komatsu A, Khosla A, Nelson DC, Kyozuka J, Flematti GR, Waters MT (2021) Desmethyl butenolides are optimal ligands for karrikin receptor proteins. New Phytol 230 (3):1003–1016. doi:10.1111/nph.17224

Zheng J, Hong K, Zeng L, Wang L, Kang S, Qu M, Dai J, Zou L, Zhu L, Tang Z, Meng X, Wang B, Hu J, Zeng D, Zhao Y, Cui P, Wang Q, Qian Q, Wang Y, Li J, Xiong G (2020) Karrikin Signaling Acts Parallel to and Additively with Strigolactone Signaling to Regulate Rice Mesocotyl Elongation in Darkness. Plant Cell 32 (9):2780–2805. doi:10.1105/tpc.20.00123

Zhou F, Lin Q, Zhu L, Ren Y, Zhou K, Shabek N, Wu F, Mao H, Dong W, Gan L, Ma W, Gao H, Chen J, Yang C, Wang D, Tan J, Zhang X, Guo X, Wang J, Jiang L, Liu X, Chen W, Chu J, Yan C, Ueno K, Ito S, Asami T, Cheng Z, Lei C, Zhai H, Wu C, Wang H, Zheng N, Wan J (2013) D14-SCF(D3)-dependent degradation of D53 regulates strigolactone signalling. Nature 504 (7480):406–410. doi:10.1038/nature12878

