## SupplementalFigures for "*Physcomitrium patens* SMXL homologs are PpMAX2-dependent negative regulators of growth": PpSMXL-SupplFiguresRev.pdf

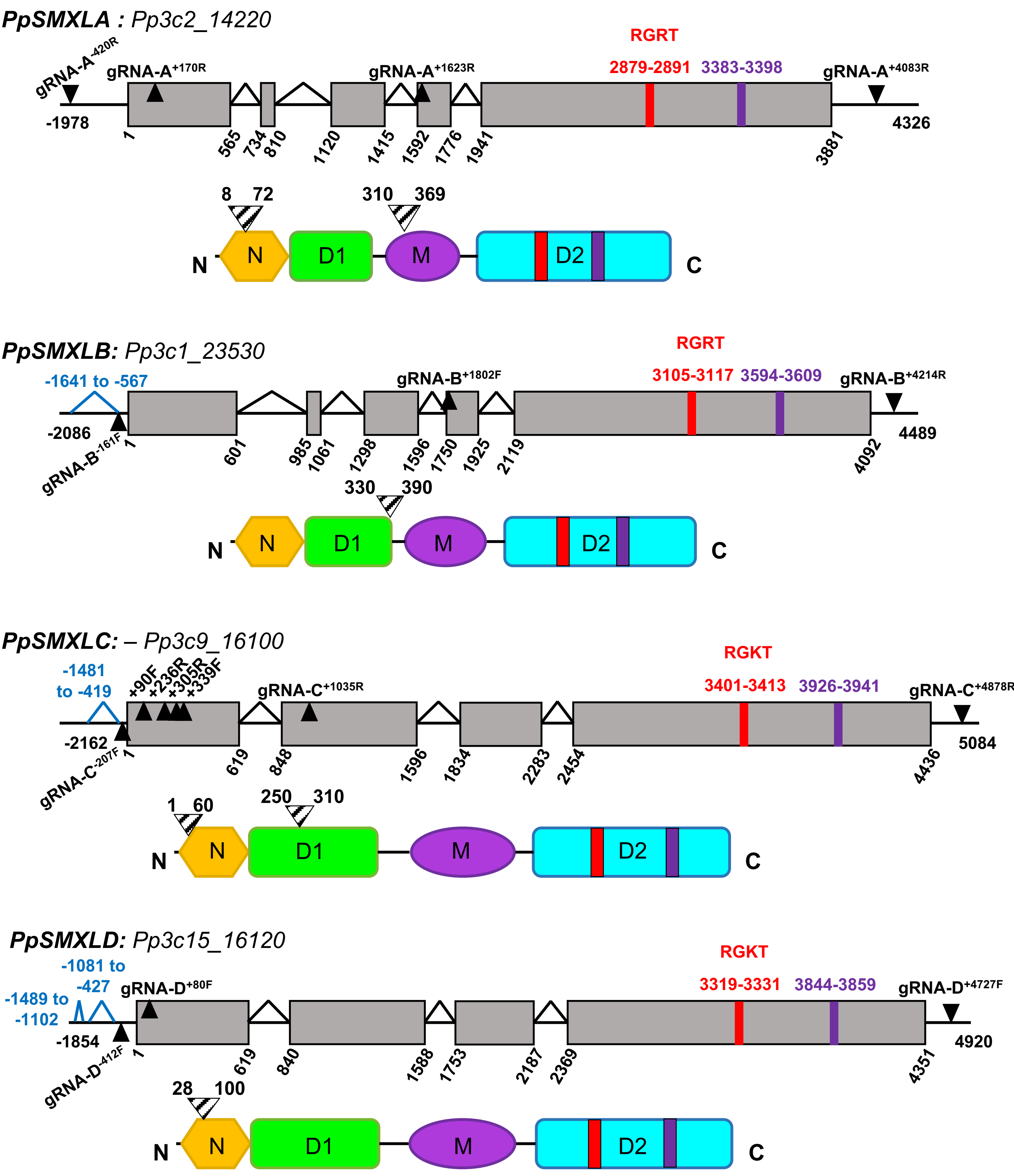


**Supplemental Figure S1 Model of *PpSMXL* genes and proteins.** Genomic sequences of the four *PpSMXL* genes were recovered from Phytozome (V3 of *P. patens* genome). Architecture of the primary (V3.1) transcript is shown for each. Only exons and inter-exonic introns are represented true to scale. Exons are shown as grey boxes, inter-exonic introns as black angles, UTRs as solid black lines and introns located in the 5'UTR are depicted as blue angles. Guide RNAs used for mutagenesis, named according to their position relative to the ATG and to their orientation, are depicted as black triangles. Positions are indicated relative to the START codon. Protein models (below gene models) are color-coded as follows: Yellow = Double ClpN domain. Green = D1 (first ATPase domain). Purple = middle domain. Cyan = D2 (second ATPase domain). Ranges of alignments shown on Supplemental Figure S4 are indicated as hatched arrow-heads. In gene and protein models, the degron motif (RGKT or RGRT) is shown as a red box and the EAR motif as a purple box. *Supports Figure 1*

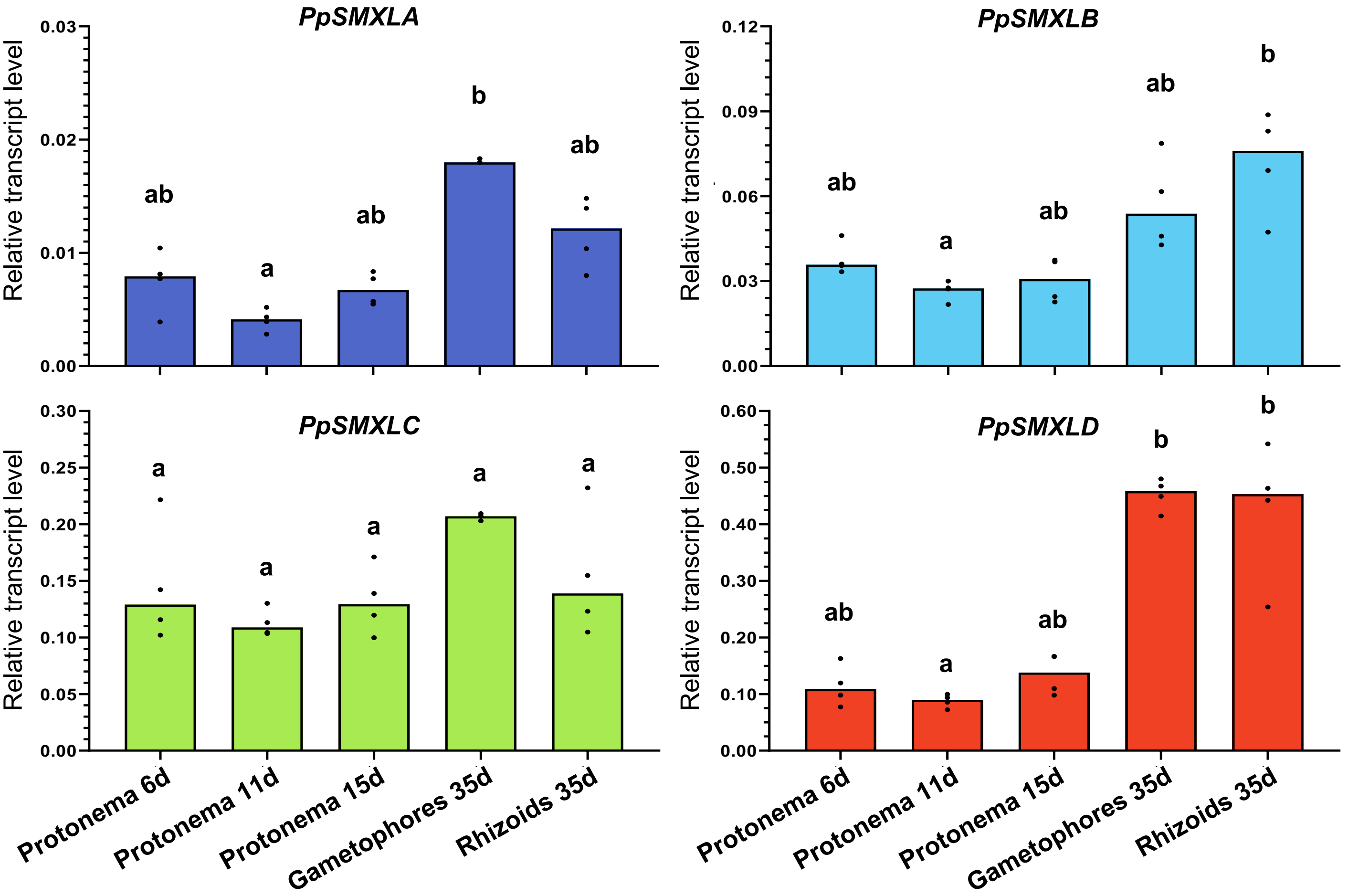


**Supplemental Figure S2 Expression of *PpSMXL* genes along *P. patens* vegetative development.** Transcript levels of the four *PpSMXL* genes, relative to the two reference genes *PpELig2* (*Pp3c14_21480*) and *PpAPT* (*Pp3c8_16590*). RT-qPCR data used for the analysis was extracted from 4 biological replicates, each with 2 technical repeats. Each point represents the mean of two technical repeats. Some points were excluded from analysis following an outliers identification test carried out in GraphPad Prism (version 8.4.2). Statistical significance scores of comparisons among tissues are indicated as bold letters (Kruskal-Wallis test followed by a Dunn *post-hoc* test. $p$<5%). Note the difference in expression values on y axes. . *Supports Figure 1*

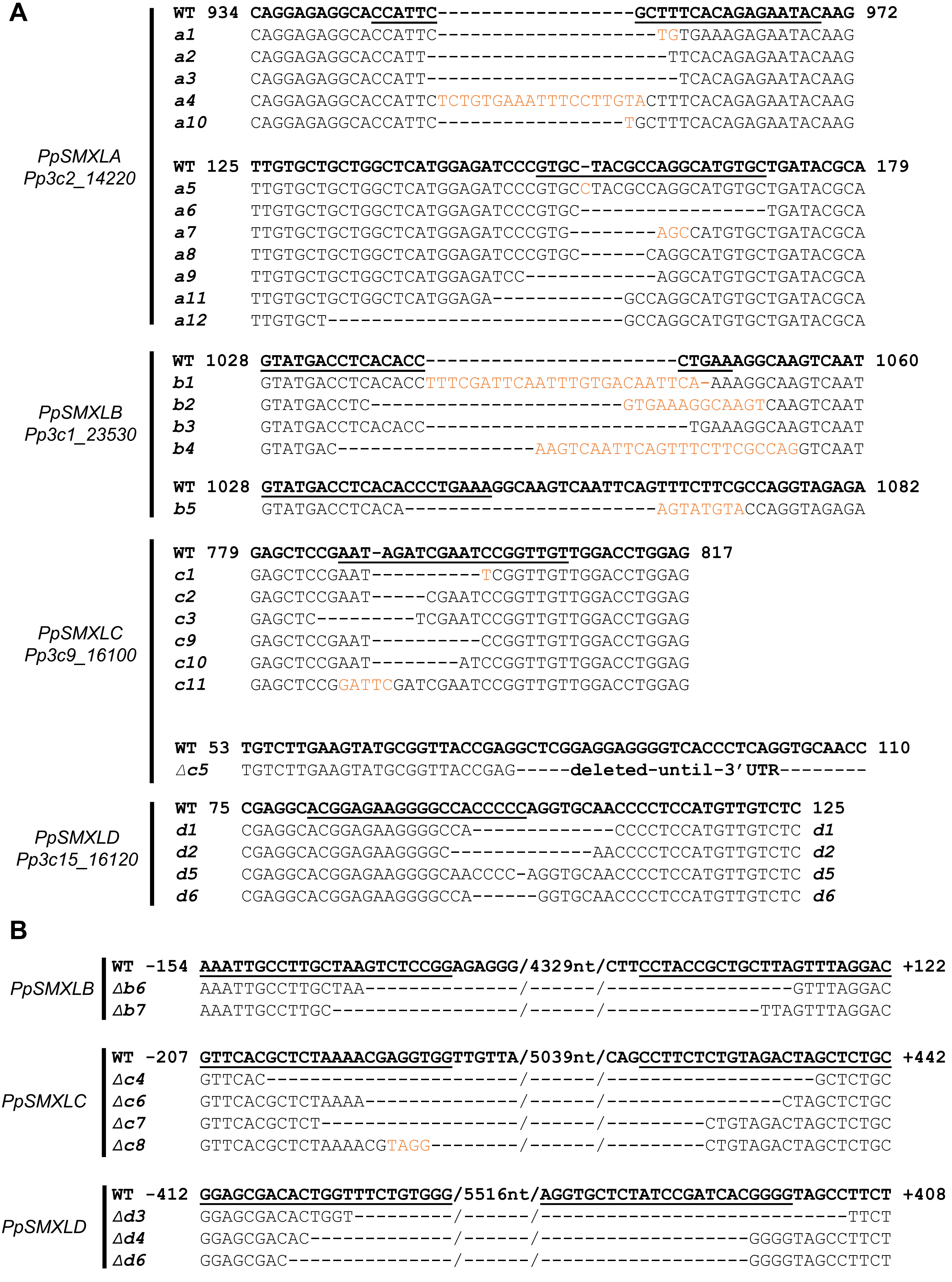


**Supplemental Figure S3 Used *Ppsmxl* and Δ*Ppsmxl* mutations.** WT sequences are given in bold and guide RNA are underlined. A, *Ppsmxl* sequences. numbers refer to the position in the CDS relative to the start codon. B, Δ*Ppsmxl* sequences, numbers refer to the position in the UTRs relative to the start and stop codons (- for the 5'UTR and + for the 3'UTR). . *Supports Figure 1 and Figure 3*

*PpSMXLA*

```
WT   310 TEQERHHSLSQRIQELRRKWQFVCSNSHSERCITMDSSPARDRGLGPPSHNWMKASILNN 369
a1   310 TEQERHHSVKENTRAPS 326
a2   310 TEQERHHFHREYKSSVVNGNLFAVTHTQKDASLWTLRQLEIGDLVRHLITG 360
a3   310 TEQERHHFTENTRAPS
a4   310 TEQERHHSL 318
a10  310 TEQERHHSAFTENTRAPS 327
```

```
WT   8 VQNTLSLPAQQVLRQAISAARERGHAQVQPLHVAFVLLAHGDPVLRQACADTHSQTLHGLHQCHA 72
a5   8 VQNTLSLPAQQVLRQAISAARERGHAQVQPLHVAFVLLAHGDPVPTPGMC 57
a6   8 VQNTLSLPAQQVLRQAISAARERGHAQVQPLHVAFVLLAHGDPVLIRILIPCTGCINATL 67
a7   8 VQNTLSLPAQQVLRQAISAARERGHAQVQPLHVAFVLLAHGDPVSHVLIRILRPCTGCINATL 70
a8   8 VQNTLSLPAQQVLRQAISAARERGHAQVQPLHVAFVLLAHGDPVPGMC 55
a9   8 VQNTLSLPAQQVLRQAISAARERGHAQVQPLHVAFVLLAHGDPGMC 53
a11  8 VQNTLSLPAQQVLRQAISAARERGHAQVQPLHVAFVLLAHGEPGMC 53
a12  8 VQNTLSLPAQQVLRQAISAARERGHAQVQPLHVAFVLPGMC 48
```

*PpSMXLB*

```
WT 330 LHSQQLYRKWLLTCMTSHPERQVNSVSSPGRDGGPSLLHHSWINASIPNNAENSGLVIPKS 390
b1 330 LHSQQLYRKWLLTCMTSHLSIQFVTIQKASQFSFFAR 366
b2 330 LHSQQLYRKWLLTCMTS 346
b3 330 LHSQQLYRKWLLTCMTSHLKGKSIQFLRQVEMEDPVCYIIAGLMLVYPIMLKTLGW 385
b4 330 LHSQQLYRKWLLTCMTSQFSFFARSIQFLRQVEMEDPVCYIIAGLMLVYPIMLKTLGW 387
b5 330 LHSQQLYRKWLLTCMTSQVCTR 351
```

*PpSMXLC*

```
WT  250 ESHERRELVCRAPNRSNPVVGPGGGLSSKDEDVLNILNIFLRPRIKNVILVGDITAANAVN 310
c1  250 ESHERRELVCRAPNS---VVGPGGGLSSKDEDVLNILNIFLRPRIKNVILVGDITAANAVN 310
c2  250 ESHERRELVCRAPNRIRLLDLEVVFHRRMRTSSTF 284
c3  250 ESHERRELVCRALESGCWTWRWSFIEG 276
c9  250 ESHERRELVCRAPN---PVVGPGGGLSSKDEDVLNILNIFLRPRIKNVILVGDITAANAVN 310
c10 250 ESHERRELVCRAPNIRLLDLEVVFHRRMRTSSTF 283
c11 250 ESHERRELVCRAPDSIESGCWTWRWSFIEG 279
```

```
WT   1 MRSGANSVQHLLTPAAHGVLKYAVTEARRRGHPQVQPLHVVSMLLTHAGSRLRQACMLSQ 60
Δc5  1 MRSGANSVQHLLTPAAHGVLKYAVTELEFKLV 32 (Alternative in-frame STOP codon)
```

*PpSMXLD*

```
WT 28 RRRGHPQVQPLHVVSMLLTHAESRLRQACMLSHPHNSQAAECRALEVCFNVALDHLPQSALAATSQPILSNAL 100
d1 28 RRRGHPSMLSPCSSLTRPAVCDKRVCYRIRTTLRPQSAGLWRFASMLHWTIFRSPRWLLPASLFCQMP 95
d2 28 RRRGNPSMLSPCSSLTRPAVCDKRVCYRIRTTLRPQSAGLWRFASMLHWTIFRSPRWLLPASLFCQMP 95
d5 28 RRRGHPRCNPSMLSPCSSLTPRAVCDKRVCYRIRTTLRPQSAGLWRFASMLHWTIFRSPRWLLPASLFCQMP 99
d6 28 RRRG--QVQPLHVVSMLLTHAESRLRQACMLSHPHNSQAAECRALEVCFNVALDHLPQSALAATSQPILSNAL 100
```

**Supplemental Figure S4 PpSMXL predicted mutant proteins local alignments.** Numbers in black indicate the range of local alignments, also placed on protein models (see Supplemental Figure S1). WT sequences are given in bold, and colored according to the predicted protein domains (see Supplemental Figure S1). Stretches of variant amino acids finishing with a premature STOP codon are written in red. Small insertions/deletions of amino acids are noted in orange. Other Δ mutations are not shown here as mutant alleles cannot generate proteins (see supplemental Figure S3 for genomic DNA alignments). *Supports Figure 1 and Figure 3*

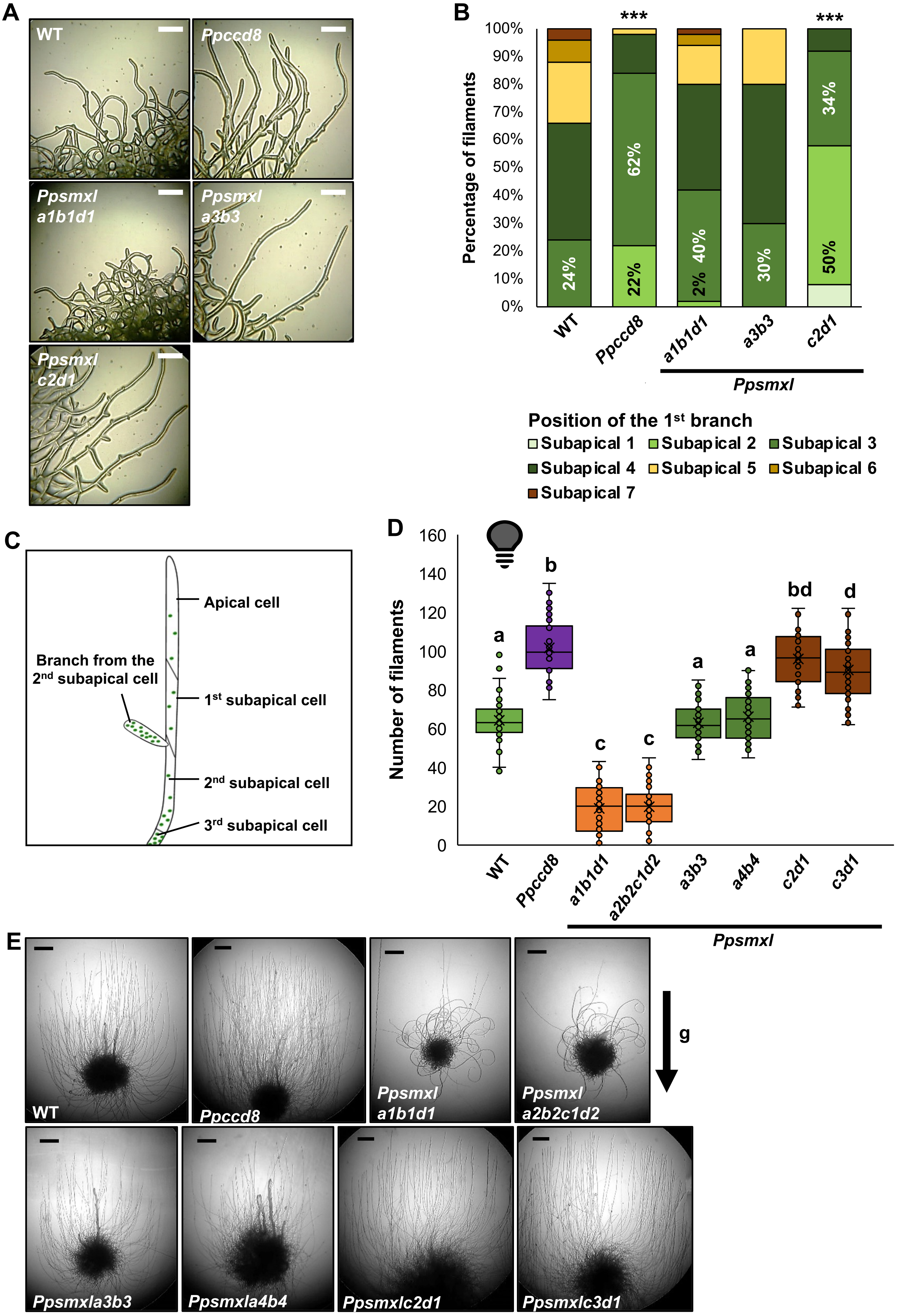

A
WT
Ppccd8
Ppsmxl a1b1d1
Ppsmxl a3b3
Ppsmxl c2d1
B
***
***
Percentage of filaments
100%
90%
80%
70%
60%
50%
40%
30%
20%
10%
0%
24%
22%
62%
2%
40%
30%
50%
34%
WT
Ppccd8
a1b1d1
a3b3
c2d1
Ppsmxl
Position of the 1st branch
Subapical 1
Subapical 2
Subapical 3
Subapical 4
Subapical 5
Subapical 6
Subapical 7
C
Apical cell
Branch from the 2nd subapical cell
1st subapical cell
2nd subapical cell
3rd subapical cell
D
Number of filaments
160
140
120
100
80
60
40
20
0
a
b
c
c
a
a
bd
d
WT
Ppccd8
a1b1d1
a2b2c1d2
a3b3
a4b4
c2d1
c3d1
Ppsmxl
E
WT
Ppccd8
Ppsmxl a1b1d1
Ppsmxl a2b2c1d2
g
Ppsmxla3b3
Ppsmxla4b4
Ppsmxlc2d1
Ppsmxlc3d1

**Supplemental Figure S5 Further characterization of *Ppsmxl* mutants**

(A-C) Protonema branching profile of *Ppsmxl* mutants. Data was obtained from 50 filaments from 5 independent plants of each genotype. The first 8 subapical cells were taken into account. A, Representative images of protonema filaments at the periphery of individual plants grown horizontally in long days conditions on low N medium on cellophane. Scale = 200 µm. B, Percentage of filaments developing their first branch at different subapical positions. Statistically significant differences in the proportion of filaments first branching from subapical cells 1 to 3 were found between WT and *Ppccd8* (***), WT and *Ppsmxlcd* (***) (two-sided Fisher's exact tests). Percentages are given on the graph for positions 2 and 3. C, Schematic diagram of a protonema filament with identification of subapical cells. (D-E) Growth of *Ppsmxl* mutants in the dark. Plants were grown in control light conditions for 10 days. then placed vertically in the dark for an additional 10 days. D, The number of caulonema filaments was counted for each plant. Statistical significance of differences between groups are given by bold letters (Welch's ANOVA ($p<0.0001$) followed by a Dunnett T3 *post-hoc* test. n=41-48 plants). E, Pictures of representative individuals at the end of the experiment. Scale bars = 2 mm. g = gravity. *Supports Figure 1*

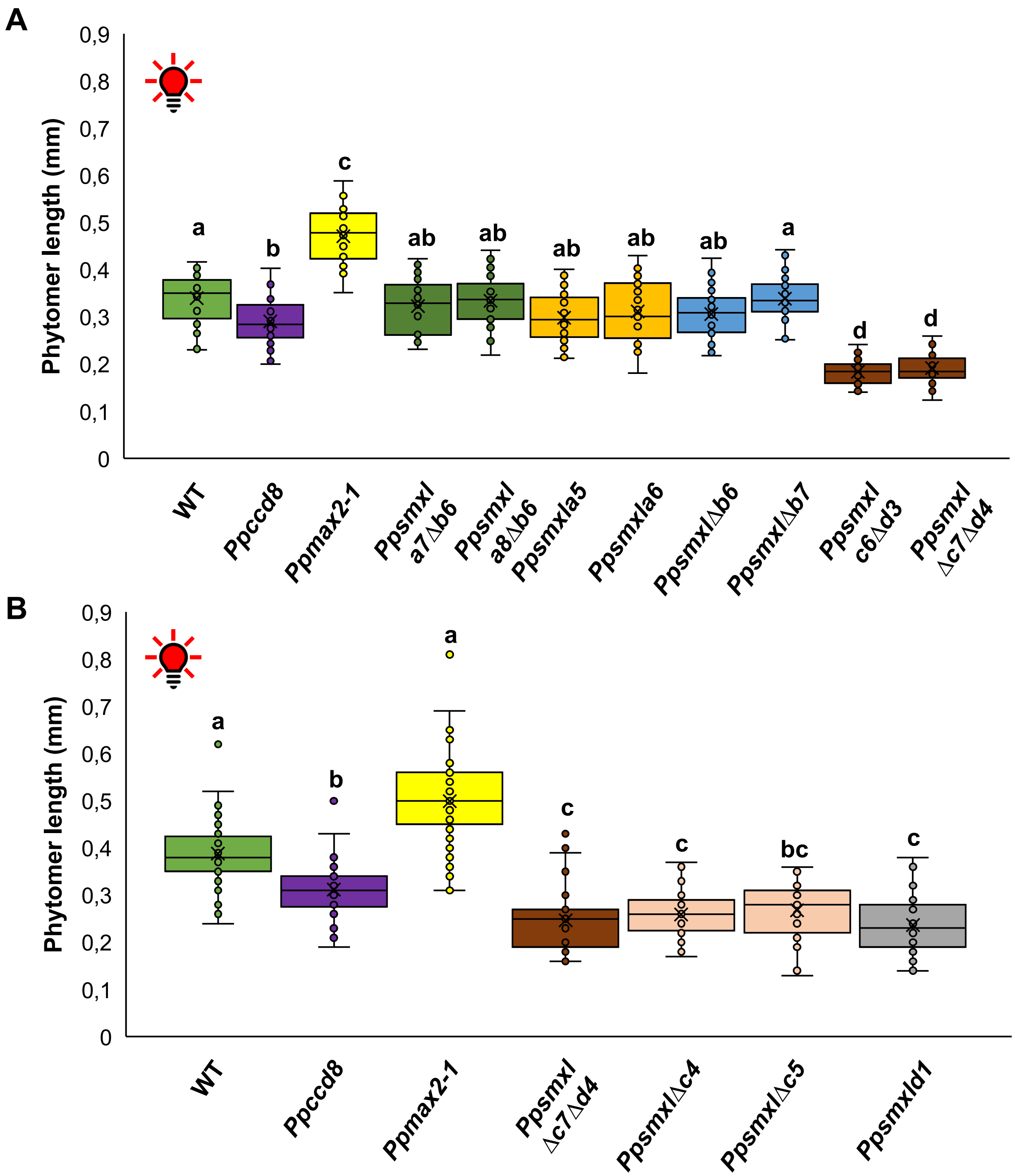


**Supplemental Figure S6 Growth of *Ppsmxl* mutant gametophores in red light.** Estimated length of phytomers of two-month-old gametophores measured in Figure 1 ($n \geq 30$ gametophores. for each genotype). Length of phytomers was estimated by dividing gametophore length by phyllid number, for each individual gametophore. A, Estimation relative to data from Figure 1E. Statistical significance of comparisons between all genotypes are indicated by bold letters (Welch's ANOVA ($p<0.0001$) followed by a Dunnett T3 *post-hoc* test. n=30). B, Estimation relative to data from Figure 1F. Statistical significance of comparisons between all genotypes are indicated by bold letters (Kruskal Wallis test ($p<0.0001$) followed by a Dunn *post-hoc* test. n=43-45). *Supports Figure 1.*

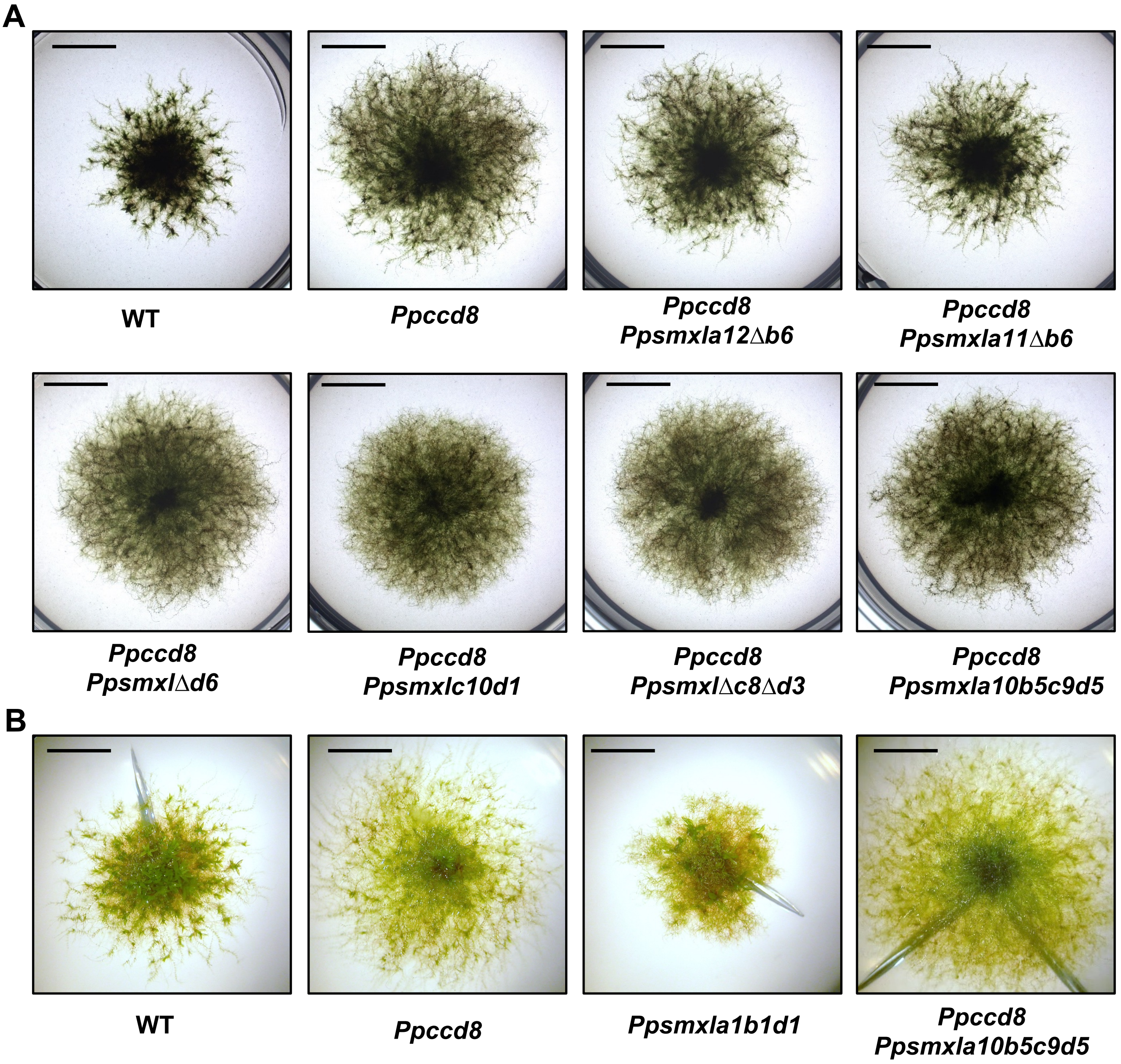


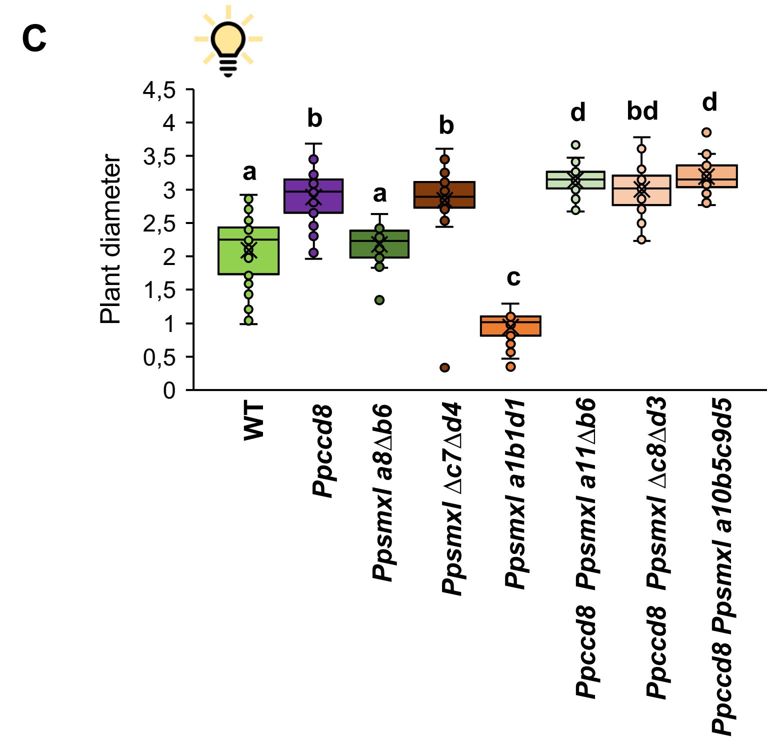


**Supplemental Figure S7 Genetic analysis of *PpSMXL* relationship with *PpCCD8***
A, Phenotype of three-week-old plants on low nitrogen content medium (without underlying cellophane). B, Phenotype of three-week-old plants on low nitrogen content medium (with underlying cellophane). Scale bar = 5 mm. C, Plant diameter of some of these mutants on low nitrogen content medium (with underlying cellophane), after 5 weeks growth. Statistical significance of comparison between all genotypes is indicated by bold letters (Welch's ANOVA ($p<0.0001$) followed by a Dunnett T3 *post-hoc* test. n =34-39). *Supports Figure 1 and 2.*

**A** *PpMAX2 – Pp3c17_1180*

```
WT      -20 TGGTGTAGCAGAGGCAAGACATGGAAGCA/39-nt/CGTCAGTCAATGCAATCTTCCCCAGAGGCGGC 80
max2-16     TGGTGTAGCAGAG----------------/-----/---------------------------GCGGC
```

**B**

```
WT      60 ACRDWYFIERQTRTELSLRGNICVMHELPTCFQQICTLDLSQCSPWGSSLFQSTQNGEEIGNCLRIGFPNV 130
max2-16    --first-83-aa-deleted---MHELPTCFQQICTLDLSQCSPWGSSLFQSTQNGEEIGNCLRIGFPNV 130
```

**C**

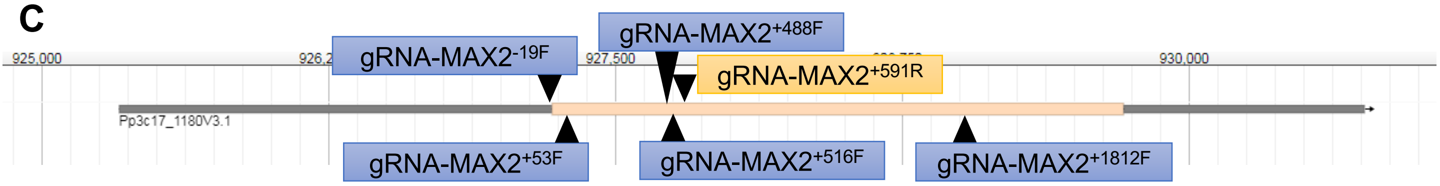


**Supplemental Figure S8 Description of *Ppmax2-16* mutation.** A, Genomic sequences. The WT sequence is given in bold green and numbers refer to the position in the *PpMAX2* gene relative to the start codon (in purple). Guide RNAs are underlined. B, Predicted protein sequences. The end of the putative F-box domain is written in bold blue. C, Location of sequences recognized by the five guide RNAs used for CRISPR-Cas9 mediated mutagenesis of *PpMAX2* (guide RNAs are named according to their location relative to the ATG and according to their orientation). *Supports Figure 3*

**A**

| Experiment | WT (–)-dGR24 1µM | *Ppccd8* (–)-dGR24 1µM | *Ppmax2-1* (–)-dGR24 1µM | *Ppsmxlab* (–)-dGR24 1µM | *Ppsmxlcd* (–)-dGR24 1µM |
|---|---|---|---|---|---|
| 1 | 43.7% | 20.6% | -3.6% | 42.8% | 11.8% |
| 2 | 11.9% | 4.8% | -1.8% | 11.9% | 15.2% |
| 3 | 24.7% | -3.9% | -37.2% | 13.4% | 30.4% |
| 4 | 37.7% | 3.8% | | 22.3% | -0.8% |
| **Mean** | **29.5%** | **6.3%** | **-14.2%** | **22.6%** | **14.2%** |
| **SEM** | **7.1%** | **5.1%** | **11.5%** | **7.1%** | **6.4%** |

**B**

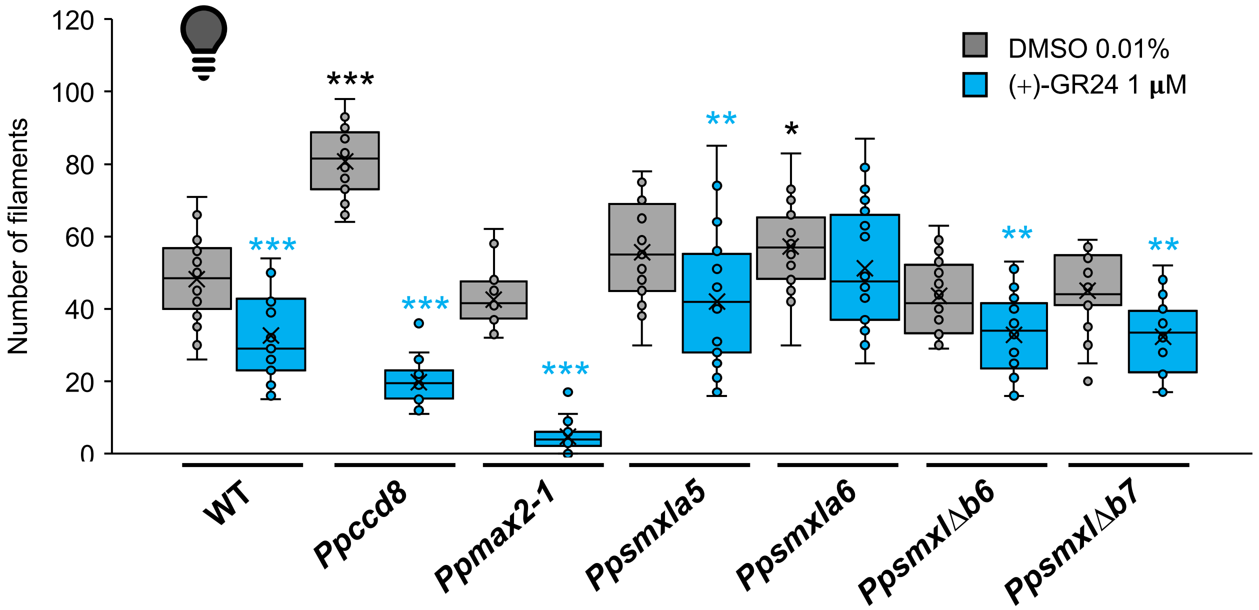


**C**

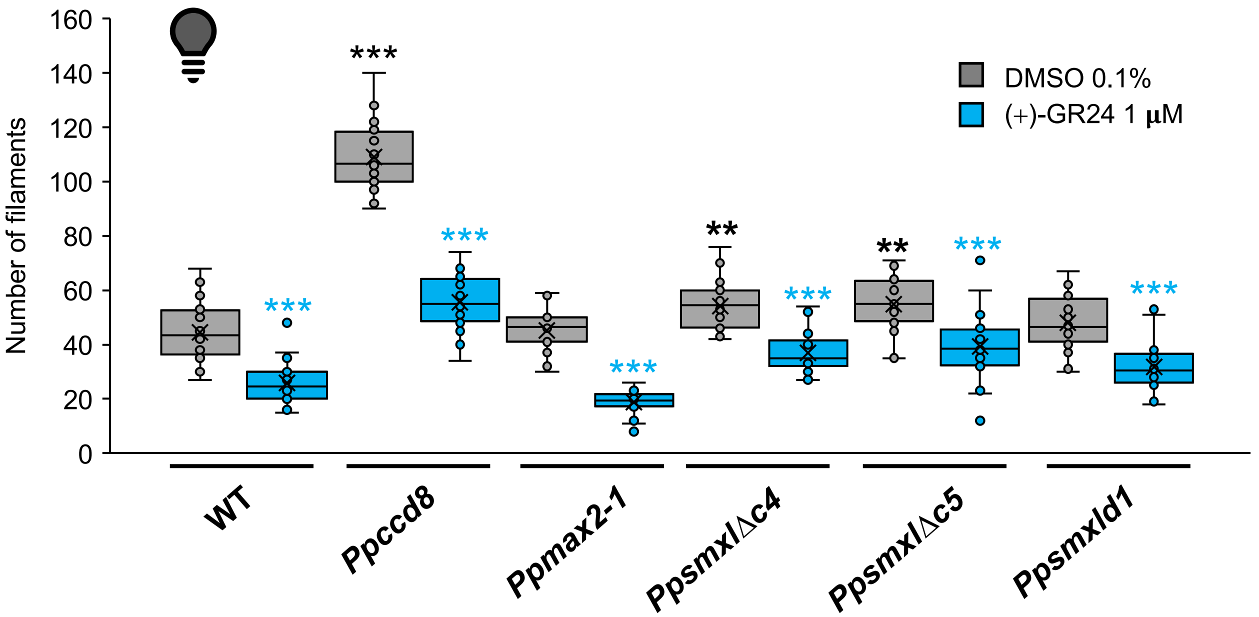


**D**

| Experiment | WT (+)-GR24 1µM | *Ppccd8* (+)-GR24 1µM | *Ppmax2-1* (+)-GR24 1µM | *Ppsmxlab* (+)-GR24 1µM | *Ppsmxlcd* (+)-GR24 1µM |
|---|---|---|---|---|---|
| 1 | -19.0% | -52.3% | -16.2% | -14.6% | -19.9% |
| 2 | -15.5% | -42.2% | -34.8% | -30.3% | 0.7% |
| 3 | -27.7% | -36.6% | -85.8% | -24.9% | -7.2% |
| 4 | -12.9% | -46.6% | | -26.9% | -7.1% |
| 5 | -32.2% | -75.6% | -89.0% | -28.7% | -2.2% |
| **Mean** | **-21.5%** | **-50.7%** | **-56.5%** | **-25.1%** | **-7.1%** |
| **SEM** | **3.7%** | **6.7%** | **18.3%** | **2.8%** | **3.5%** |

**Supplemental Figure S9 Phenotypic response of *P. patens* to (+)-GR24 and (−)-desmethyl-GR24 in the dark.** A, Response of WT, *Ppccd8, Ppmax2-1* and double mutants *Pp*smxl*a7*Δ*b6* and *Ppsmxl*Δ*c6*Δ*d3* to 1µM of (−)-desmethyl-GR24 ((−)-dGR24) in the dark. Response ratios obtained in three to four independent experiments are shown. Data shown in Figure 4A and B are those from experiment#1. B-D, Phenotypic response of *Ppsmxl* single mutants to (+)-GR24 in the dark. B,C: two-week-old plants of each genotype were mock treated with 0.01% DMSO (grey) or 1 µM of (+)-GR24 (blue). Plants were incubated vertically in the dark for ten days then negatively gravitropic *caulonema* filaments were enumerated for each plant. B, Statistical significance of comparisons of control groups relative to WT is shown as bold black symbols (standard ANOVA ($p<0.0001$) followed by a Dunnett *post-hoc* test). Statistical significance of comparisons between control and treated for each genotype is shown as bold blue symbols (two-tailed Student t-tests, except for P*pmax2-1* for which a two-tailed Mann-Whitney test was carried out). C, Statistical significance of comparisons of control groups relative to WT is shown as bold black symbols (standard ANOVA ($p<0.0001$) followed by a Dunnett *post-hoc* test). Statistical significance of comparisons between control and treated for each genotype is shown as bold blue symbols (two-tailed Student t-tests). For all statistical analyses, p-values are reported as * $0.01\leq p<0.05$. ** $0.001\leq p<0.01$. *** $p<0.001$. D, Response of WT, *Ppccd8, Ppmax2-1* and double mutants *Pp*smxl*a7*Δ*b6* and *Ppsmxl*Δ*c6*Δ*d3* to 1 µM of (+)-GR24 in the dark. Results of 4 to 5 independent experiments are shown, where the gain or loss of filaments response has been calculated for each (+)-GR24-treated group as the following ratio: ($number_{treated}$ - $number_{mock}$) / $number_{mock}$. Data shown in Figure 4C are those from experiment#5. *Supports Figure 4.*

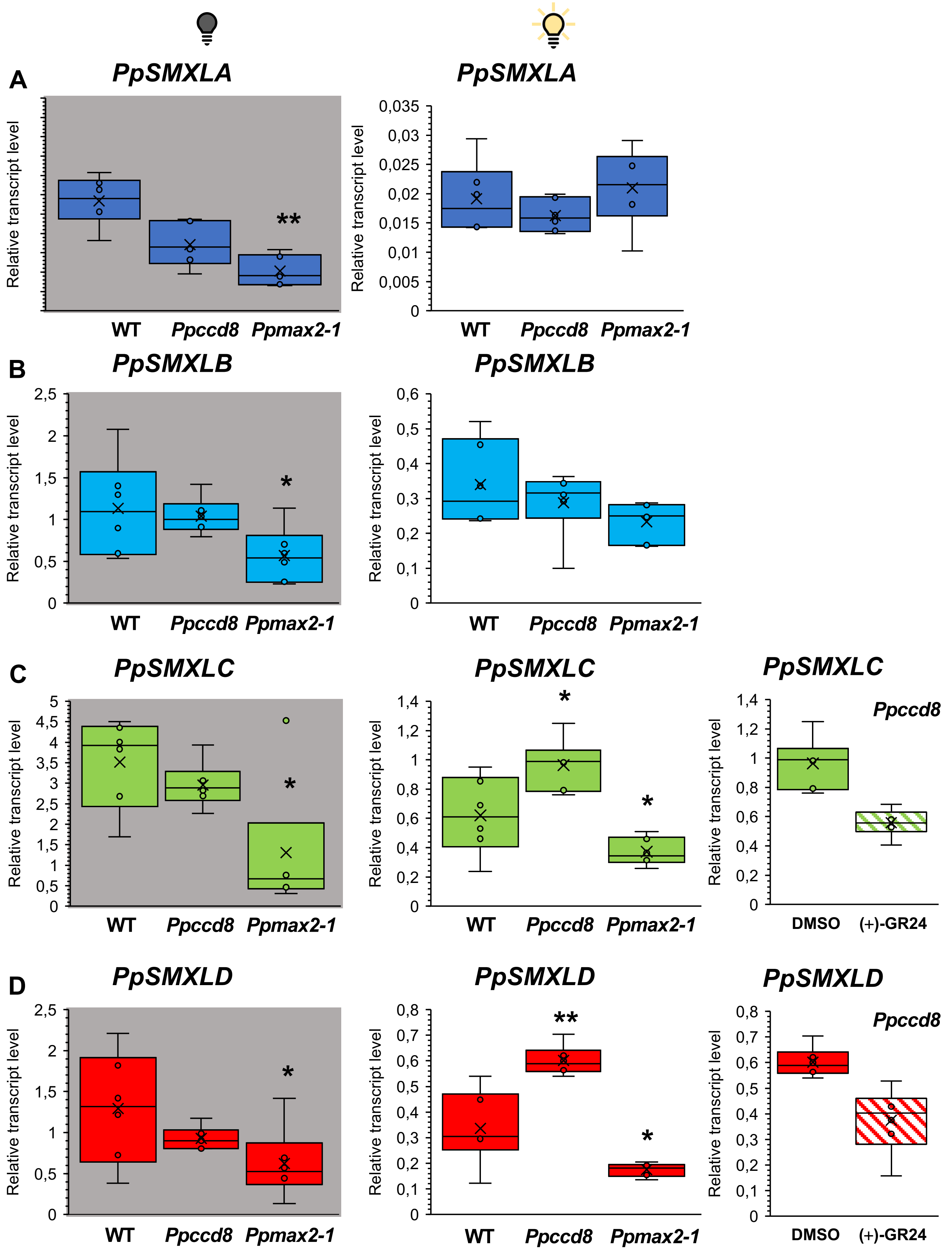


**Supplemental Figure S10 Expression of *PpSMXL* genes in response to light, in WT, *Ppccd8* and *Ppmax2-1*.** Transcript levels of the four *PpSMXL* genes, relative to the two reference genes *PpElig2* (*Pp3c14_21480*) and *PpAPT* (*Pp3c8_16590*). Two-week-old plants (n = 6 pools of plants) were incubated in the dark for one week (left) or kept in usual long-day conditions with white light illumination (middle panel). Right panels: *Ppccd8* mutant plants grown in light were treated with 1 µM (+)-GR24 (hatched plots) or DMSO (control, filled plots) for 6 hours. Stars indicate confidence level of a one-tailed Mann-Whitney test between WT and mutant (p<5%). *Supports Figure 1 and Figure 8*

```
AtSMXL7  (85-117) : RLPSSKSTPTTTVEED--PPVSNSLMAAIKRSQAT
AtSMXL6  (85-114) : RLPSSKS-PAT--EED--PPVSNSLMAAIKRSQAN
PpSMXLA (83-113) : RLQQCSSSGS-TVNLLGL---SNALVAALKRAQAQ
PpSMXLB (83-112) : RLPQSSSSS--TVHPLGL---SNALVAALKRAQTH
PpSMXLC (82-110) : HLPQSALAAASQP---IL---SNALMAALKRAHAH
PpSMXLD (82-110) : HLPQSALAATSQP---IL---SNALMAALKRAHAH
OsSMAX1 (99-131) : RLPAAAAAAAAHGAGASPPVSNALVAALKRAQAQ
```

**Supplemental Figure S11 Alignment of the region characterized as a functional NLS in rice SMAX1 (Choi *et al.* 2020). Arabidopsis SMXL7 (Liang *et al.* 2016). and *P. patens*.** Numbers in parentheses indicate the range of aligned sequences (in amino acids). Residues that are conserved between all seven SMXL are in red, those that are conserved in at least two SMXL (relative to AtSMXL7) are in blue. *Supports Figure 5*

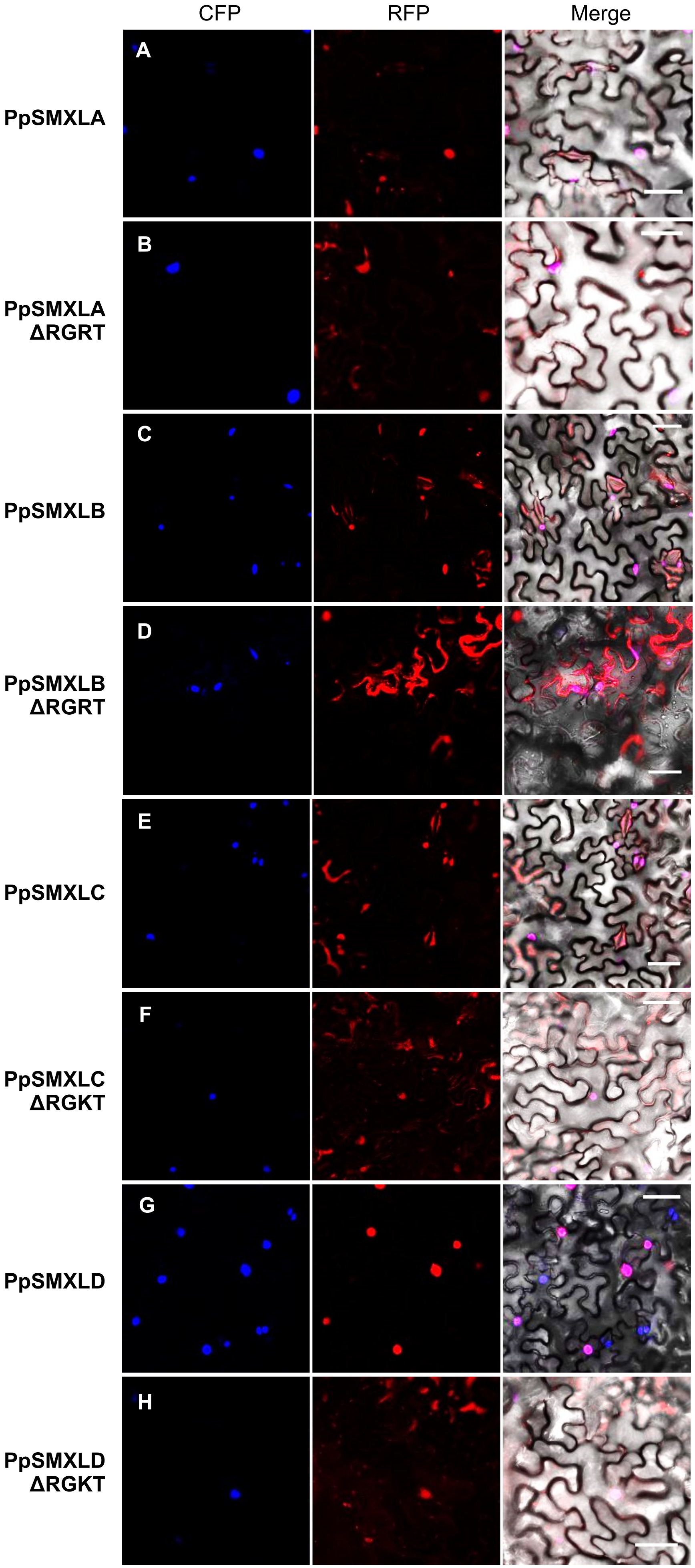


**I**

| | |
|---|---|
| **Walker A consensus** | **XGXXXXGXGKTXXXX** |
| AtSMAX1 | **GDGNSSFRGKTALDK** |
| OsSMAX1 | **DGPNMGFWGKTALDR** |
| PsSMAX1 | **DSDAHHIRGKTVLDR** |
| AtSMXL6 | **CSLDDKFRGKTVVDY** |
| AtSMXL7 | **DSLDDRFRGKTVVDY** |
| OsD53/OsD53L | **DWDDSSFRGKTGIDC** |
| PpSMXLA | **EIDGMQYRGRTAVDS** |
| PpSMXLB | **EIDGLQLRGRTAEDS** |
| PpSMXLC | **DDSGMRYRGKTPLDR** |
| PpSMXLD | **ETDDFRMRGKTPLDR** |

**Supplemental Figure S12 Subcellular localization of RFP-PpSMXL fusion proteins in *Nicotiana benthamiana* leaves. and effect of P-loop deletion on RFP-PpSMXL fusion proteins stability and localization.** A-H, Transient expression of RFP tagged PpSMXL proteins (red) was carried in a *N. benthamiana* line stably expressing H2b-CFP (blue), to achieve high expression of *RFP-PpSMXLA* (A), *RFP-PpSMXLA-ΔRGRT* (B), *RFP-PpSMXLB* (C), *RFP-PpSMXLB-ΔRGRT* (D), *RFP-PpSMXLC* (E), *RFP-PpSMXLC-ΔRGKT* (F), *RFP-PpSMXLD* (G) and *RFP-PpSMXLD-ΔRGKT* (H). I, Local alignment of the Walker A/P-loop « degron » motif of SMXL proteins from Arabidopsis (At), rice (Os), pea (Ps) and *P. patens* (Pp). Residues that are conserved between all included SMXL are in red. *Supports Figure 5*

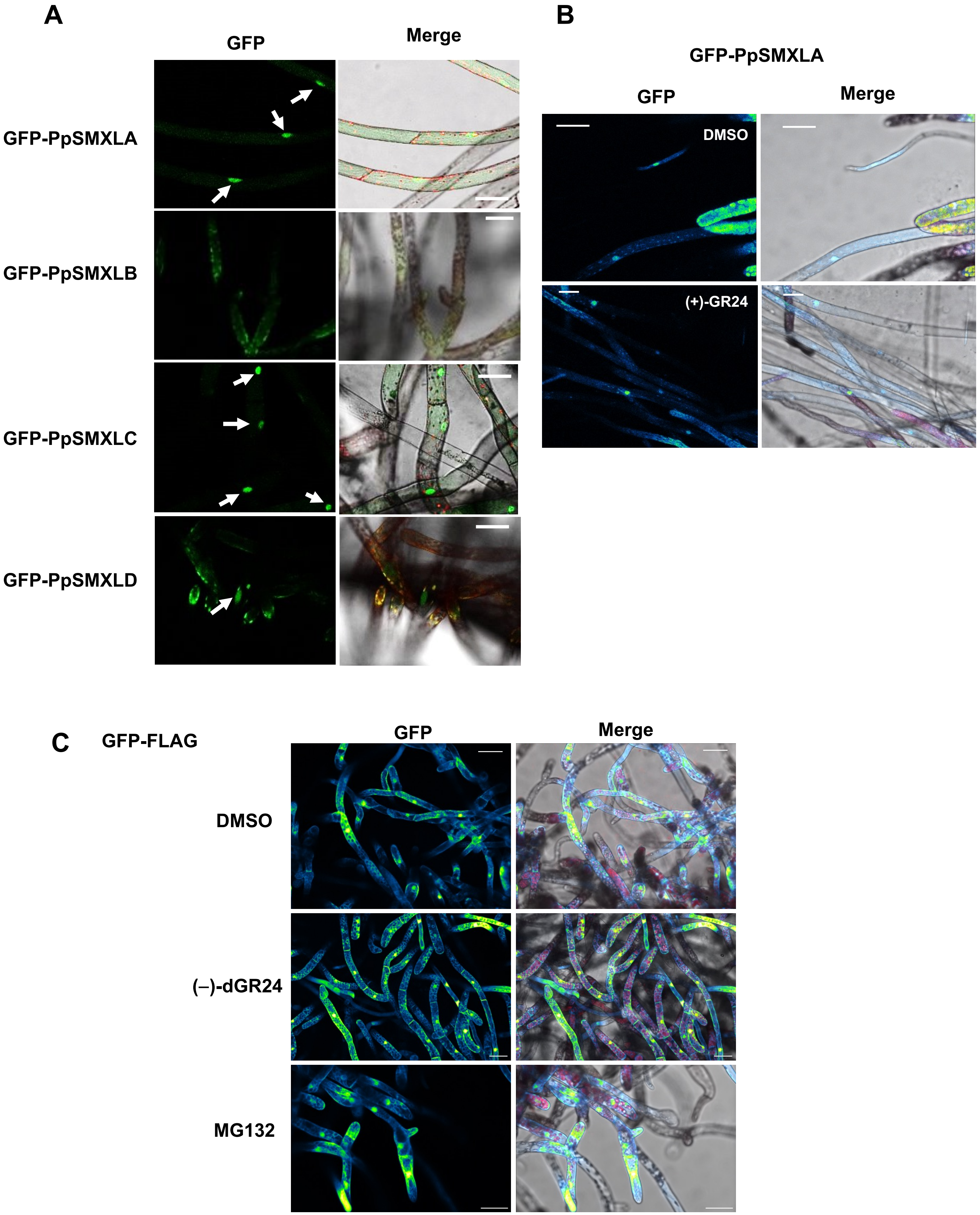


**Supplemental Figure S13 Subcellular localization of GFP-PpSMXL fusion proteins in *P. patens*.** A, Confocal images of protonema filaments of transgenic *P. patens* lines expressing *proZmUbi:GFP-PpSMXL* fusions as indicated on the left. Merge are overlays of autofluorescence (red) and GFP (green) and bright field channels. Bar = 50 µm. Arrow points to a nucleus. B-C Effect of a the SL mimic (+)-GR24 treatment on OE-SMXLA (A) or GFP-FLAG control line (B). Filaments were incubated in a solution of DMSO 0.01% (control) or in (+)-GR24 10 µM (diluted in 0.01% DMSO) for 20 minutes, or in MG132 100 µM (diluted in 0.01% DMSO) for 20 minutes. Merge are overlays of GFP (Green Fire Blue LUT), autofluorescence (Red) and bright field images. Scale bar = 50 µm. *Supports Figure 5*

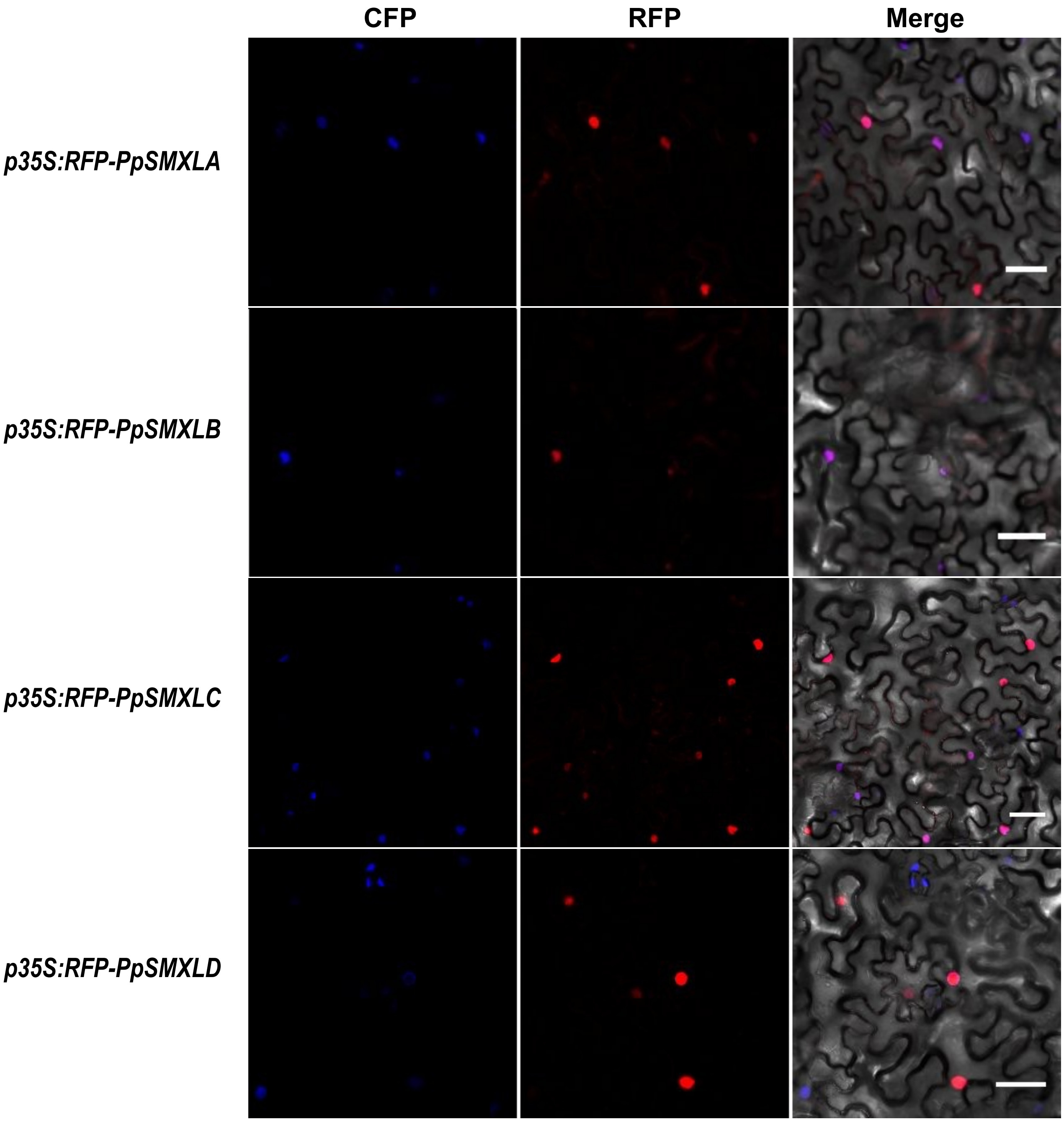


**Supplemental Figure S14 Subcellular localization of RFP-PpSMXL fusion proteins in *Nicotiana benthamiana* leaves in response to a (+)-GR24 treatment.** Infiltrations were carried out in a *N. benthamiana* line stably expressing H2b-CFP. Leaf pieces were immerged in a 5µM (+)-GR24 solution (diluted in 0.1% DMSO) for 20 minutes before observation. Merge are overlays of CFP (blue), RFP (red) and bright field images. Bar = 50 µm. *Supports Figure 5 and Supplemental Figure 13*

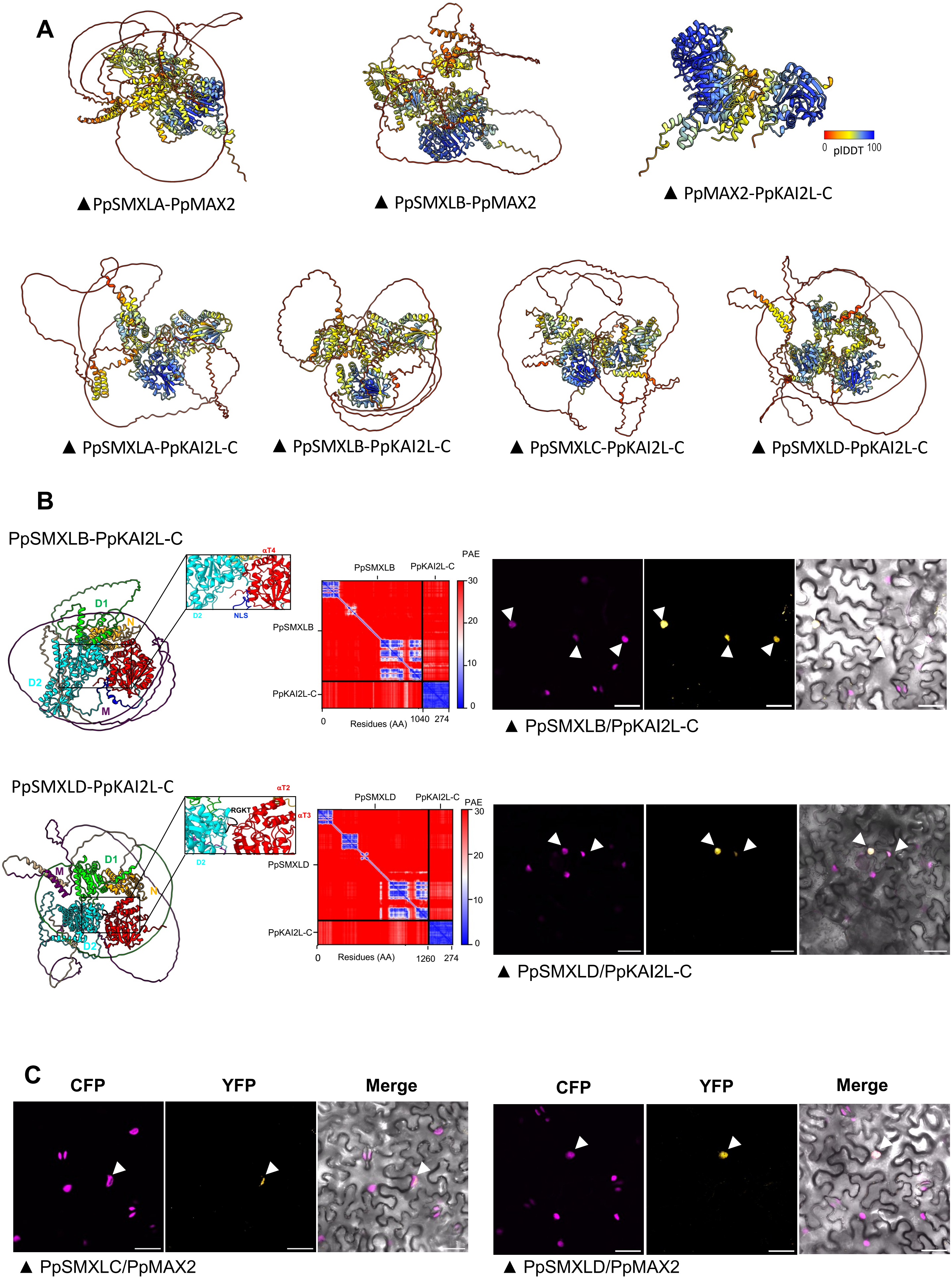
A
PpSMXLA-PpMAX2
PpSMXLB-PpMAX2
PpMAX2-PpKAI2L-C
0 pIDDT 100
PpSMXLA-PpKAI2L-C
PpSMXLB-PpKAI2L-C
PpSMXLC-PpKAI2L-C
PpSMXLD-PpKAI2L-C
B
PpSMXLB-PpKAI2L-C
αT4
D1
N
D2
NLS
M
PpSMXLB
PpKAI2L-C
PAE
Residues (AA)
PpSMXLB/PpKAI2L-C
PpSMXLD-PpKAI2L-C
αT2
αT3
RGKT
PpSMXLD
PpSMXLD/PpKAI2L-C
C
CFP
YFP
Merge
PpSMXLC/PpMAX2
PpSMXLD/PpMAX2

**Supplemental Figure S15 PpSMXL protein interaction predictions and BiFC assays in *Nicotiana benthamiana*.** See also Figure 6. A, ColabFold models of PpSMXLA-PpMAX2, PpSMXLB-PpMAX2, PpMAX2-PpKAI2L-C, PpSMXLA-PpKAI2L-C, PpSMXLB-PpKAI2L-C, PpSMXLC-PpKAI2L-C, and PpSMXLD-PpKAI2L-C interactions, as indicated below each model, and colored by per-residue model confidence score (pLDDT). High and low pLDDT values indicate strong and low confidence in the predicted structure, respectively. B, ColabFold models of, from top to bottom, PpSMXLB-PpKAI2L-C and PpSMXLD-PpKAI2L-C (AlphaFold). The PpSMXL proteins are colored by domains as used in Supplemental Figure S1, with N in orange, D1 in lime, M in purple and D2 in cyan. PpKAI2L-C is represented in red and the four alpha helices of the lid domain are annotated as αT1, αT2, αT3 and αT4 respectively. Predicted Aligned Error (PAE) values for the models are shown next to the models. Low PAE values indicate strong confidence in the distances between two amino acids, and high values indicate low confidence. Insets: Interaction domains highlighted for every generated model: for PpSMXLB-PpKAI2L-C, a NLS (Nuclear Localization Signal) is predicted in the D2 domain and colored in dark blue, for PpSMXLD-PpKAI2L-C the RGKT (degron) motif is colored in black. C, BiFC assays in *N. benthamiana* show that PpSMXLC and PpSMXLD may interact with PpMAX2. Below each tryptic of images, the first indicated protein is fused to the N-terminal part of eYFP, while the second protein is fused to the C-terminal part (both tags are fused at the N-terminal end of *P. patens* proteins). Colocalization of CFP-H2b and eYFP biFC signals are pointed at with white arrowheads. Merge are overlays of CFP (magenta). YFP (yellow hot LUT) and bright field images. Bar = 50 µm. Negative controls are shown in Supplemental Figure S16. *Supports Figure 6*

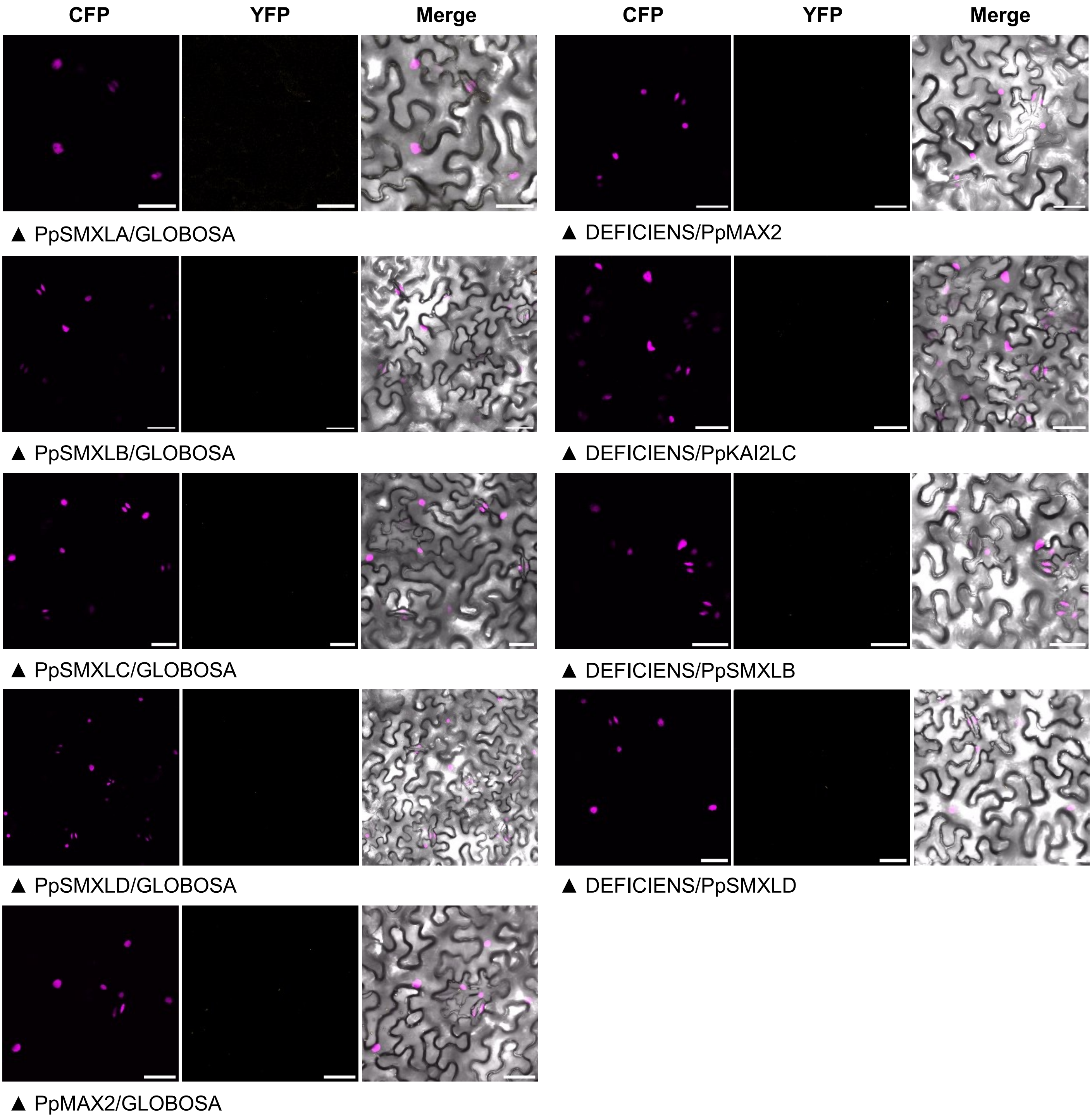


**Supplemental Figure S16 Controls for BiFC assays shown Figure 6 and Supplemental Figure S14**
Tested interactions with DEFICIENS and GLOBOSA proteins from *Antirrhinum major* (negative controls). Below each tryptic of images. the first indicated protein is fused to the N-terminal part of eYFP. while the second protein is fused to the C-terminal part (both tags are fused at the N-terminal end of *P. patens* proteins). Merge are overlays of CFP (magenta). YFP (yellow hot LUT) and bright field images. Bar = 50 µm. The previously published GLOBOSA/DEFICIENS interaction (Azimzadeh et al.. 2008) was consistently used a positive control of eYFP reconstruction. *Supports Figure 6 and Supplemental Figure S15*

A *pSMXL5*

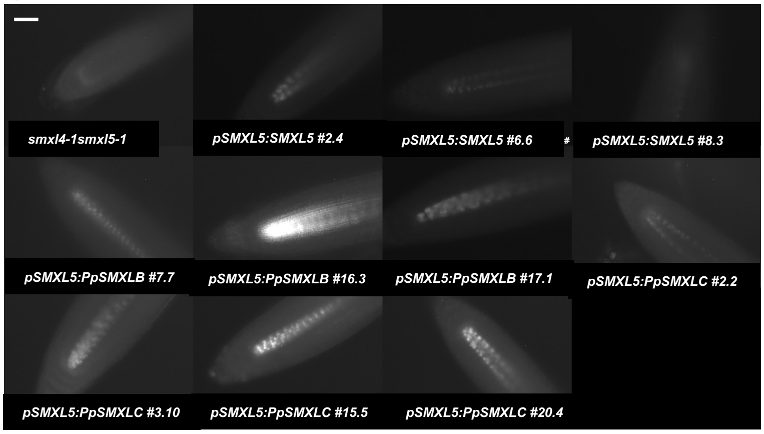


B *pSMAX1*

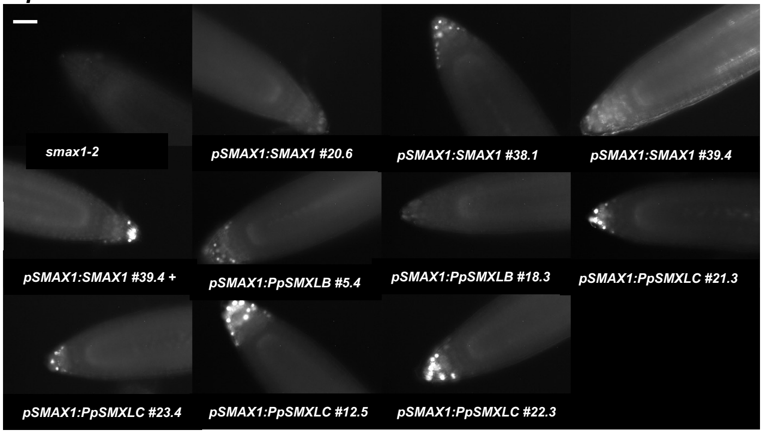


**Supplemental Figure S17**

**A-B m-Citrine epifluorescence in transformed Arabidopsis seedling root tips.** A. *smxl4.5* (*smxl4-1.smxl5-1*) double mutant (control. top left) was transformed with constructs carrying the indicated CDS fused to m-Citrine. under the control of SMXL5 (pSMXL5) promoter. B. *smax1-2* mutant (control. top left) was transformed with constructs carrying the indicated CDS under the control of SMAX1 promoter. For *pSMAX1:SMAX1 #39.4* +. the seedling has been pre-treated with 100 µM MG132. Bar = 30 µm.

**C. Semi Quantitative RT-PCR** on rosette leaf cDNA from Arabidopsis Col0. and transformed Arabidopsis plants as indicated. using primers specific for Arabidopsis (*EF1α. AtSMXL6*) or *P. patens (PpSMXLB. PpSMXLC*) genes (see Supplemental Table 1). *P. patens:* cDNA from 3 week-old moss plants. *Supports Figure 7 and Supplemental Figure S18.*

C

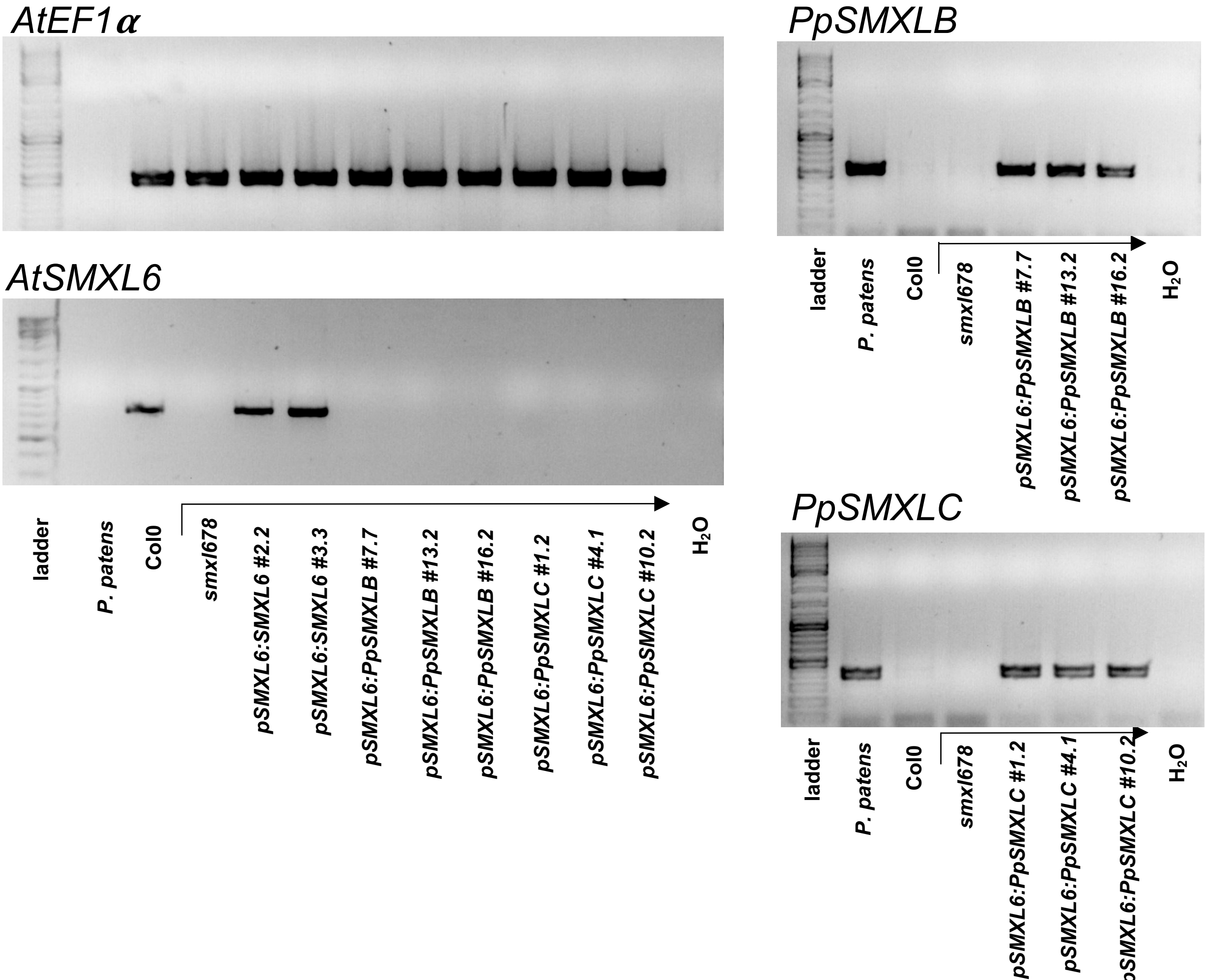

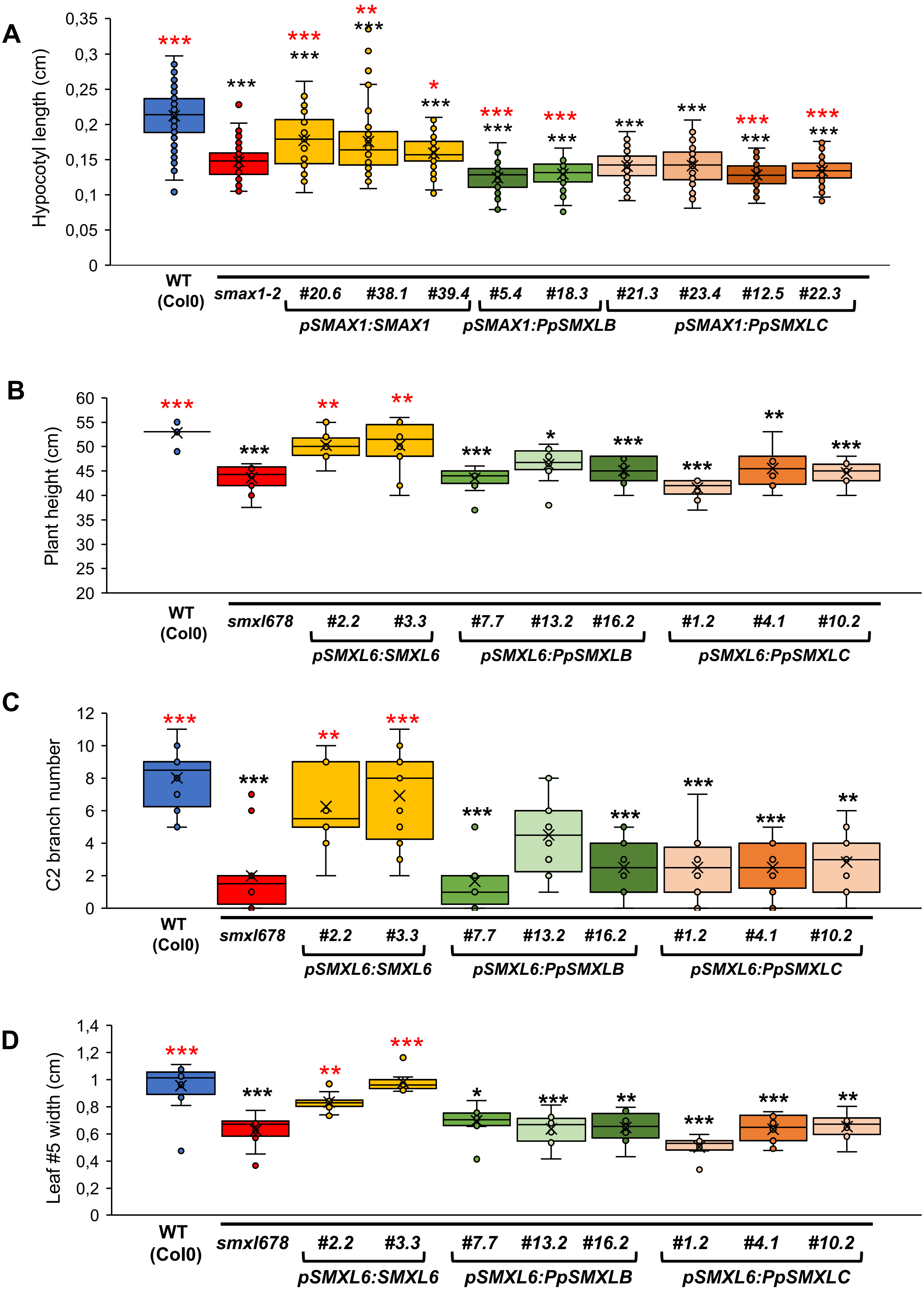
A
Hypocotyl length (cm)
0,35
0,3
0,25
0,2
0,15
0,1
0,05
0
WT (Col0)
smax1-2
#20.6
#38.1
#39.4
#5.4
#18.3
#21.3
#23.4
#12.5
#22.3
pSMAX1:SMAX1
pSMAX1:PpSMXLB
pSMAX1:PpSMXLC
B
Plant height (cm)
60
55
50
45
40
35
30
25
20
WT (Col0)
smxl678
#2.2
#3.3
#7.7
#13.2
#16.2
#1.2
#4.1
#10.2
pSMXL6:SMXL6
pSMXL6:PpSMXLB
pSMXL6:PpSMXLC
C
C2 branch number
12
10
8
6
4
2
0
WT (Col0)
smxl678
#2.2
#3.3
#7.7
#13.2
#16.2
#1.2
#4.1
#10.2
pSMXL6:SMXL6
pSMXL6:PpSMXLB
pSMXL6:PpSMXLC
D
Leaf #5 width (cm)
1,4
1,2
1
0,8
0,6
0,4
0,2
0
WT (Col0)
smxl678
#2.2
#3.3
#7.7
#13.2
#16.2
#1.2
#4.1
#10.2
pSMXL6:SMXL6
pSMXL6:PpSMXLB
pSMXL6:PpSMXLC

**Supplemental Figure S18 Arabidopsis complementation assays.** A. Transformants in the *smax1-2* background. Hypocotyl length of 11-day-old seedlings grown under low light. For control lines (Col0 and *smaxl1-2*). n=89-129 seedlings per genotype. grown on 5-7 different plates. For all transformed lines. n≥ 45 seedlings per genotype. grown on at least 3 different plates. Statistical significance of comparisons between WT (Col0) and each genotype is shown as black symbols and statistical significance of comparisons between *smax1-2* and each genotype is shown as red symbols (for both: Welch ANOVA (p<0.0001) followed by a Dunnett T3 *post-hoc* test. n = 45-129); B-D. Transformants in the *Atsmxl678* background. B. Height of 5-week-old plants; C. Caulinary (C2) branch number of 5-week-old plants; D. Width of the 5$^{th}$ leaf of 3-week-old plants. B-D**.** Statistical significance of comparisons between WT (Col0) and each genotype is shown as black symbols. and statistical significance of comparisons between *smxl678* and each genotype is shown as red symbols (Kruskal-Wallis tests (p<0.0001) followed by Dunn *post-hoc* tests. n=12). *Supports Figure 7*
